# Dendritic modulation enables multitask representation learning in hierarchical sensory processing pathways

**DOI:** 10.1101/2022.11.25.517941

**Authors:** Willem A.M. Wybo, Matthias C. Tsai, Viet Anh Khoa Tran, Bernd Illing, Jakob Jordan, Abigail Morrison, Walter Senn

## Abstract

While sensory representations in the brain depend on context, it remains unclear how such modulations are implemented at the biophysical level, and how processing layers further in the hierarchy can extract useful features for each possible contextual state. Here, we first demonstrate that thin dendritic branches are well suited to implementing contextual modulation of feedforward processing. Such neuron-specific modulations exploit prior knowledge, encoded in stable feedforward weights, to achieve transfer learning across contexts. In a network of biophysically realistic neuron models with context-independent feedforward weights, we show that modulatory inputs to thin dendrites can solve linearly non-separable learning problems with a Hebbian, error-modulated learning rule. Finally, we demonstrate that local prediction of whether representations originate either from different inputs, or from different contextual modulations of the same input, results in representation learning of hierarchical feedforward weights across processing layers that accommodate a multitude of contexts.

## Introduction

Sensory processing in the brain is commonly thought of as proceeding through an increasingly abstract and invariant hierarchy of representations^1,2^. According to this view, neurons have a fixed tuning to specific stimuli: in early sensory areas neurons identify basic features such as lines, gratings^3^ or simple auditory waveforms^4^, whilst neurons further in the processing stream are selective to faces^5,6^, speakers^7^ or words^8^. Artificial neurons in feedforward network models also exhibit such receptive field properties, and similarity between responses in these networks and in sensory brain regions lends support to this view of sensory processing^9,10^. However, the activity of sensory neurons is not driven purely by bottom-up inputs, but is also modulated by internal mental states^11^. These modulating inputs, relayed by top-down connections from various cortical areas (Fig 1A), communicate high-level information about behavioural context^12–14^, task demands^15–17^, expectations^18–20^, motor commands^19,21,22^ and memory^23,24^.

**Figure 1.**
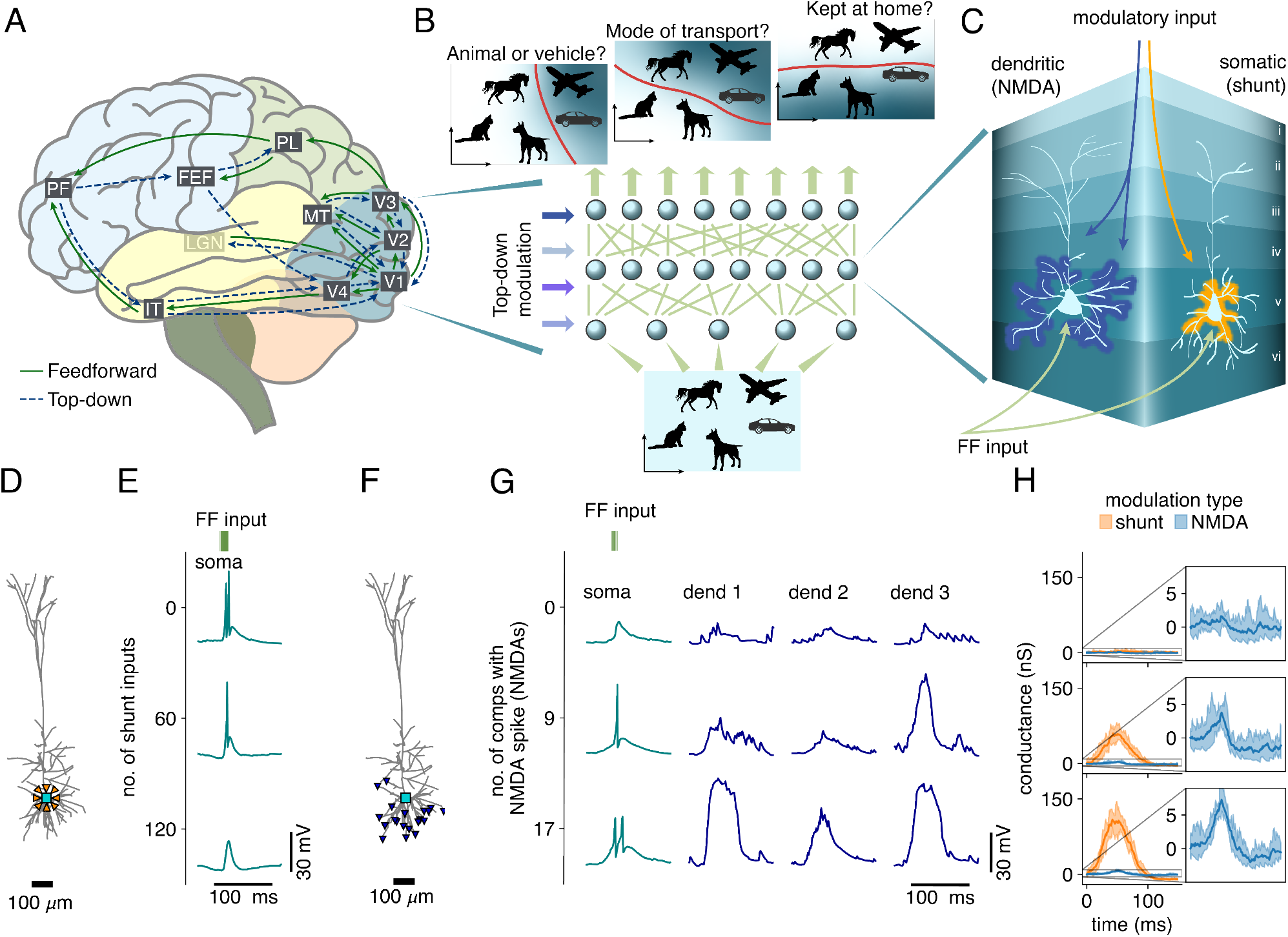
Contextual modulation of neurons in sensory processing pathways. **A:** Top-down connections from prefrontal and motor areas relay high-level information to early sensory processing neurons (adapted from Gilbert et al.^11^). **B:** We hypothesize that high-level information from prefrontal and motor areas modulates the activity of early sensory neurons, enhancing response properties of neurons with task-relevant receptive fields. These modulations induce a task-dependent functional remapping of sensory processing pathways built on fixed, task-agnostic feedforward connectivity. **C:** At the biophysical level, we investigate two plausible candidate mechanisms that could implement quasi-tonic neuron-specific modulations: somatic shunting inhibition and dendritic NMDA spikes. **D:** L5 PC model configuration to investigate somatic shunting: feedforward and shunting (orange) inputs target the somatic compartment (teal). **E:** The somatic response to identical feedforward inputs (top, green, Gaussian burst of 175 inputs), for three modulation levels resulting in zero, one or two output spikes. **F:** L5 PC model configuration to investigate dendritic modulation: modulatory inputs target up to 20 dendritic compartments (blue), whereas feedforward inputs target the somatic compartment (teal). **G:** Somatic responses (left, teal) to identical feedforward inputs (top, green, Gaussian burst of 40 inputs), for three different levels of dendritic modulation (right, blue) resulting in zero, one or two output spikes. **H:** Comparison of effective conductance changes, as measured at the soma, between shunt and NMDA modulation for the three modulation levels shown in E and G.

While it is attractive to assume that such top-down connections to sensory areas adapt feedforward processing to the many contexts that may occur in natural environments, the computational utility of modulating neurons at all levels in the processing stream remains poorly understood. Such modulations induce a dependence on the contextual state in sensory representations at any given processing layer. Consequently, the next processing layer in the hierarchy has to be connected in such a way that it can extract useful features, not only for each possible sensory input, but also for each possible contextual state. Most artificial neural network approaches that seek to implement multitask learning avoid this complication by defining separate output networks for each task, on top of a common trunk that generates a context-independent representation of the inputs^25,26^. Nevertheless, the pervasiveness of contextual modulation in sensory processing indicates that this adaptation is an important component of cortical computation, and reshapes the functional mapping of sensory processing pathways (Fig 1B)^27^. While some authors have explored modulations to early processing layers^28–30^, their networks were trained through error backpropagation in a purely supervised fashion. Unsupervised, representation-based learning is considered more biologically plausible^31–33^, but has not been applied to context-modulated representations.

Biophysically, the way in which modulations to sensory neurons are implemented remains unknown. A probable constraint is that contextual modulations have a longer time-scale than rapid feedforward processing, where volleys of action potentials propagate upwards through the processing hierarchy^34,35^, their trajectories modulated by the contextual inputs. Two candidate mechanisms appear well adapted to implement neuron-specific modulations with long time scales. One possibility is that fast-spiking inhibitory interneurons target the somata of pyramidal cells (PCs)^36^, effectively opening quasi-tonic conductances in the membrane that decrease input resistance and render it harder for the neuron to emit action potentials (Fig 1C)^37,38^. Another possibility is that modulating inputs target thin dendritic branches^30,39,40^, opening N-Methyl-D-Aspartate (NMDA) channels and eliciting NMDA-spikes^41,42^. These events, whose duration of 50-100 ms outlasts the duration of action potentials by one to two orders of magnitude^43,44^, can also implement a sustained modulation of the neuronal output (Fig 1C).

Here, we study the modulation of feedforward processing in networks of biophysically realistic neurons. By assessing effective membrane conductance changes, we find that NMDA-spikes are the more likely candidate to modulate the neuronal input-output (IO) relation. We then study the computational features of neuron-specific modulations in abstract feedforward network models, and show that these modulations allow networks without task-specific readout components to solve multiple tasks. We find that feedforward weights that extract useful information from modulated layers can indeed be learned, because multitask performance increases with network depth. This in turn allows the network to learn new tasks by adapting solely the modulating synapses, and inspired us to ask whether unsupervised learning principles exist for feedforward weights that support multitask learning through neuron-specific modulations. While the contextual modulations in abstract models are trained through gradient descent on a classification loss, we show that our approach translates to biologically realistic spiking models equipped with a Hebbian, error-modulated learning rule for the contextual synapses. Finally, we show that context-modulated representations promote self-supervised learning across a hierarchy of processing layers, by providing a form of data augmentation for contrastive learning that allows deeper processing layers to extract general, high-level features, without the need for error backpropagation across layers. Thus, instead of being a complication, such modulations could constitute an integral feature of cortical learning.

## Results

### Biophysical implementation of neuron-specific modulations

To understand how sustained modulations of individual neurons can be implemented in cortical networks, we simulate modulatory afferents of varying strengths to a biophysically realistic layer 5 (L5) PC model^45^. We compare two possible candidate mechanisms: one where fast-spiking interneurons target the peri-somatic region of PCs, providing shunting inhibition, and another where excitatory afferents target the dendritic tips of the basal and apical oblique dendrites, eliciting NMDA-spikes (Fig 1C).

In both cases, we implement feedforward inputs as short Gaussian bursts, with a width of 6 ms, and examine conditions where, for identical bursts of feedforward input, the modulatory afferents change the number of somatic outputs between zero and two spikes. Since the first mechanism is inhibitory, we tune the number of feedforward inputs per burst so that two output spikes are emitted without modulatory input (175 feedforward inputs), and increase the number of shunt inputs until all output spikes are prevented (Fig 1D, E). Conversely, as the second mechanism is excitatory, we tune the number of feedforward inputs per burst so that no output spikes are emitted without modulatory input (40 feedforward inputs), and increase the number of inputs eliciting dendritic NMDA-spikes until two output spikes are emitted (Fig 1F, G).

An experimentally testable measure that distinguishes between the candidate mechanisms is the change in effective conductance of the neuron. The time course of this conductance can be measured in voltage clamp by repeating the same input pattern at different holding potentials, and is given by the slope of the current-voltage relationship at all time points^46^. Experimental studies estimate effective conductance changes of 1 to 10 nS^46,47^. In the case of somatic modulations through shunting inhibition, our simulations show that the effective conductance change required to modulate the output firing from two to zero spikes is between 100 and 150 nS, values far outside the experimentally measured range (Fig 1H). Conversely, the effective conductance change for modulating output firing from zero to two spikes with dendritic NMDA-spikes is between 1 and 10 nS. This demonstrates that dendritic NMDA-spikes are a biologically plausible candidate to implement neuron-specific modulations (Fig 1H), on which we will focus in the remainder of this work.

### Neuron-specific modulations as bias and/or gain changes

Conceptually, neuron-specific modulations can be thought of as changing the slope and/or threshold of the neuronal IO relationship. In abstract neuron models of the form

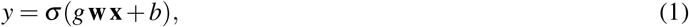

this can be implemented through modulations of gain *g* and bias *b*, with *g* primarily affecting the slope and *b* exclusively affecting the threshold. Here, *y* represents the neuronal activation, *σ* the activation function, **w** the feedforward weight vector, and **x** the feedforward input vector. Note that although *y* typically stands for the average neuronal firing rate, here we interpret it rather as the average number of somatic output spikes in response to a short burst of feedforward inputs. In this case, the ReLU activation function *σ* (*x*) = max(*x*, 0) is a reasonable choice (Fig 2A,C).

**Figure 2.**
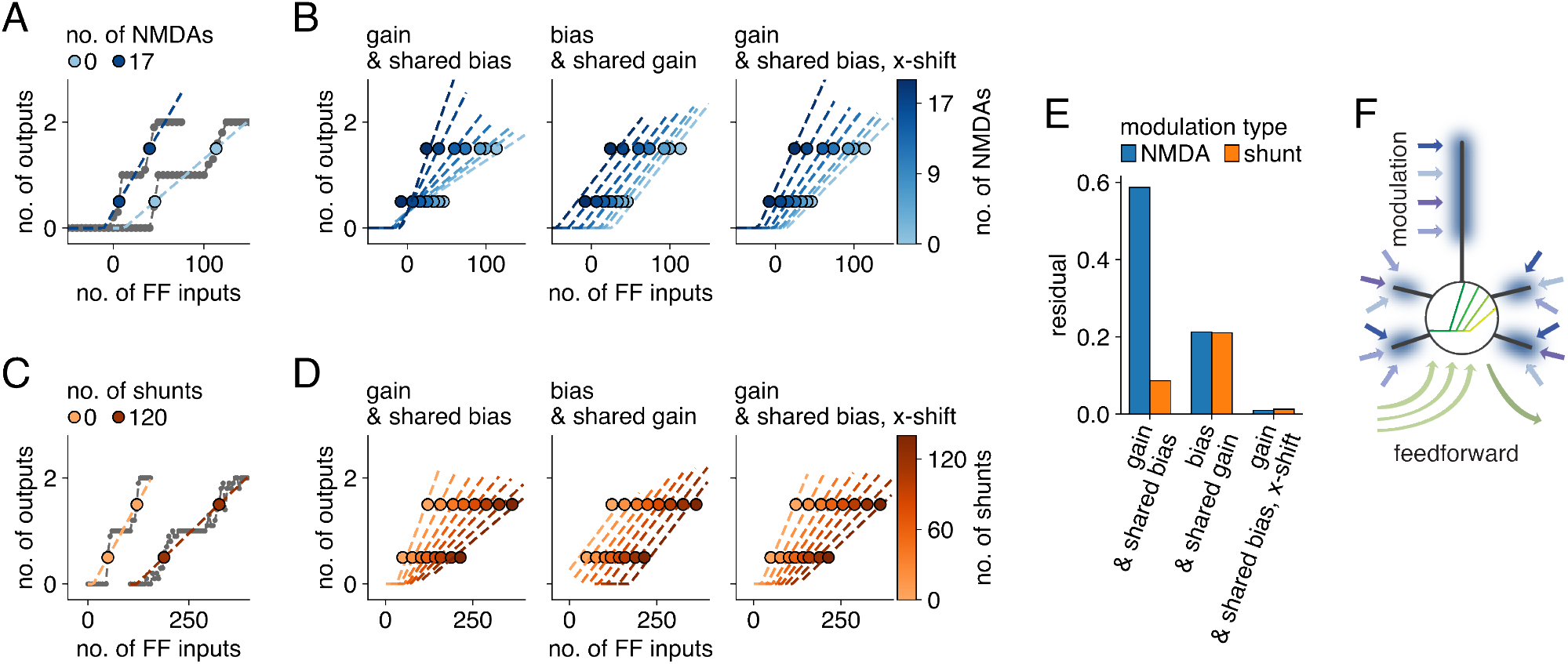
Conceptualizing neuron-specific modulations in abstract neuron models. **A:** Number of output spikes, averaged over ten trials, for two example levels of modulation (determined by the number of compartments with NMDA spikes). The x-axis shows the number of excitatory feedforward inputs (*>* 0) or inhibitory feedforward inputs (*<* 0). The thresholds (blue) where the number of emitted spikes increases are taken as the points where the linear interpolation crosses the mid-point between discrete values. We use these thresholds as fit points for the ReLU characterizing the neuronal IO relationship (dashed lines show these fits, performed here for each modulation level separately for illustrative purposes). **B:** ReLU fits to obtained threshold values (as explained in B) for eight modulation levels with a curve-specific gain and shared bias (left), a curve-specific bias and shared gain (middle) and a curve-specific gain, shared bias and additionally a shared x-shift (right). Note that two of the modulation levels had nearly overlapping thresholds, so that only seven levels are visible. **C, D:** Same as A, B, but for modulation through somatic shunting. **E:** Residual summed over the eight modulation levels for the three cases shown in B (blue) and D (orange). **F:** Proposed conceptual model of a neuron participating in sensory feedforward processing: perisomatic feedforward inputs (green) are modulated by dendritic subunits (blue), resulting in a concerted change of slope and threshold of the neuronal IO curve.

We maintain the same input configuration to the L5 PC model as before (Fig 1F), and construct IO curves for different levels of modulation by varying the number of feedforward inputs. We then model the effect of modulation on the IO dependency either as gain or bias adaptation. To fit these curves, we retain the thresholds – computed as the points where the interpolation line crosses the mid-point between discrete values – as the fit points (Fig 2A). We then fit all obtained curves together, either with a curve-specific gain and shared bias (Fig 2B, left) or with a curve-specific bias and shared gain (Fig 2B, middle). We found that the bias-modulated fits were more accurate than the gain-modulated fits (Fig 2E), but the large residual indicated that there nonetheless appeared to be a significant change in IO slope. We therefore introduced a constant *x*_shift_ parameter

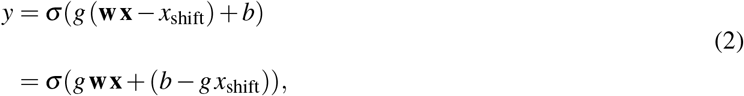

resulting in concerted additive and multiplicative modulation by gain changes. This fit produced the most accurate representation of the modulatory effect (Fig 2B, right, Fig 2E). Together, these considerations suggest a conceptual picture of sensory neurons where peri-somatic feed-forward inputs are modulated by top-down inputs impinging onto dendritic subunits (Fig 2F). These modulatory inputs increase IO slope and decrease IO threshold. For completeness, we note that somatic modulation through shunting inhibition is better fitted by pure gain modulation than bias modulation (Fig 2C-E, configuration as in Fig 1D), in agreement with prior work^37^, and that introducing a constant x-shift parameter also decreased the residual markedly.

### Multitask learning with task-dependent modulations to individual neurons

In feedforward neural network architectures, implementing task switching by providing neuron-specific modulations to the neurons in the hidden layers (Fig 3A) is a departure from the standard approach, in which task-specific output units are trained on top of a shared trunk network^25,26^. We therefore first assess whether multitask learning in this manner is even computationally feasible, and learn task-specific gains to the individual neurons in feedforward networks together with feedforward weights, x-shifts, and biases that are shared across tasks. All parameters are optimized through supervised error backpropagation.

**Figure 3.**
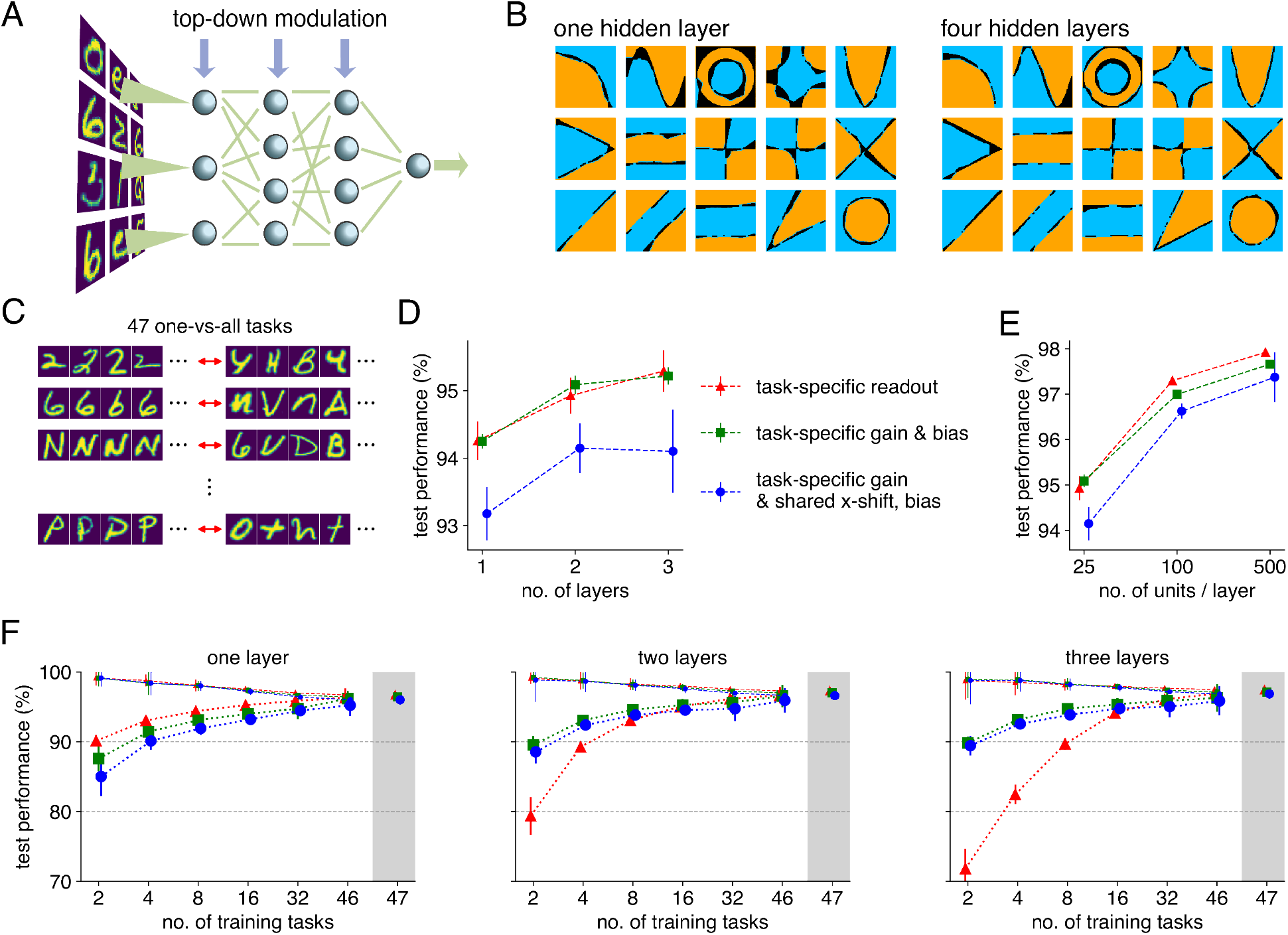
Adaptation of feedforward processing through neuron-specific modulations. **A:** Our network setup learns task-specific gains to each hidden neuron, together with a common x-shift, common bias and common feedforward weights, to implement large sets of tasks. **B:** 15 (out of 48, Fig S1A) exemplars of a two-dimensional classification multitask dataset solved with neuron-specific modulations. Correctly classifed samples are plotted in blue and orange, while incorrectly classifed samples are plotted in black. The network architecture contains one hidden layer (left) or four hidden layers (right) with 50 neurons per layer, followed by a single output unit. **C:** EMNIST is converted to a multitask learning problem by defining a one-vs-all classification task for every class in the original dataset. **D:** Performance of networks with a task-specific readout and no neuron-specific modulations (red, triangle), independently learnable task-specific gain and bias (green, square) and task-specific gain together with a shared x-shift and bias (blue, circle) as a function of the number of hidden layers (25 neurons per layer). Performance is measured by averaging over all tasks, and by additionally averaging over five initialization seeds (error bars show standard deviation of task-performance across seeds, averaged over all tasks). **E**: Same as in D but for a varying number of units per layer in networks with two hidden layers. **F**: Transfer learning performance. Networks are multitask trained on a subset of tasks (dashed line, small marker size), and performance is evaluated by training task-specific parameters on the other tasks while freezing shared parameters (averages and standard deviations computed across 128 seeds, dotted line, large marker size). Each hidden layer consists of 100 units, and multitask performances of the equivalent architecture on all tasks are shown on the right. Colors and markers as in D.

To demonstrate that neuron-specific modulations can successfully change the functional mapping of feedforward processing pathways, we train networks with one or four hidden layers to solve 48 binary classification tasks on two-dimensional inputs. These networks, each with a single set of feedforward weights, but task- and neuron-specific gains, solved all 48 tasks, demonstrating that such modulations achieve multitask learning (Fig 3B, S1A). The deeper network was more accurate (less black area in Fig 3B, S1A), indicating that multilayer architectures with neuron-specific modulations are computationally useful.

To more thoroughly test neuron-specific modulations on a dataset that is both sufficiently rich in tasks and sufficiently simple to subsequently combine with biophysical models, we convert the EMNIST dataset^48^ into a multitask learning problem (multitask EMNIST) by defining a one-vs-all classification task for every class in the original dataset (47 tasks, Fig 3C). We find that implementing neuron-specific modulations through independent gain and bias changes achieves the same performance as a task-specific readout, and that combined gain and bias changes through a constant x-shift result in a slightly reduced performance (Fig 3D, E). Qualitatively, the same behaviour is observed for both investigated forms of neuron-specific modulations: performance increases with network depth (Fig 3D), and performance increases strongly with layer size (Fig 3E). Hyperparameters, such as learning rates, are optimized for each method and architecture separately (Fig S1B). Note that we have also implemented other neuron-specific modulations Fig S1C), but the minute differences between modulation types could not be decoupled fully from choices such as network architecture, task design and training method.

In the brain, mounting evidence suggests that top-down inputs dynamically select salient features from a stable feedforward connectivity^24^. We therefore test whether our framework with neuron-specific modulations is efficient at making use of prior knowledge, encoded in the learned feedforward weights. We train shared parameters on a subset of the 47 tasks, and learn the remaining tasks by exclusively adapting the task-specific parameters. For networks with one hidden layer, we found that all approaches achieve similar transfer learning. For networks with more than one hidden layer, our approach transferred much better to the remaining tasks than networks with task-specific readouts (Fig 3F). Presuming that with more hidden layers, networks become increasingly adept at filtering out task-irrelevant information, we hypothesize that task-specific readouts for new tasks have no access to information that was not relevant for the original tasks. Conversely, neuron-specific modulations to early layers could recover such information, leading to improved transfer learning.

### Unsupervised weight matrices for networks with neuron-specific modulations

So far our supervised results have demonstrated that a network with a single set of feedforward weights, and contextual modulations to individual neurons, can solve many tasks. However, much of the learning in the brain is thought to proceed in an unsupervised fashion^49,50^. While unsupervised learning has been studied thoroughly in combination with a supervised readout on the hidden representation^32,51^, it has yet to be combined with neuron-specific modulations. We therefor investigate how to find unsupervised feedforward weight matrices that facilitate the construction of task-specific decision boundaries through supervised learning of the neuronal gains.

To explain our approach, we note that the decision of any given neuron in the feedforward pathway to become active represents a decision boundary on the sensory input space. Locally, this boundary is characterized by its normal vector (methods), which captures the input features that the neuron uses to make a decision about whether to become active, and is always a linear combination of the input weight vectors to the network (Fig 4A). A necessary condition to be able to construct a given decision boundary is that its normal vectors can all be constructed with the feedforward weight matrices (Fig 4B). Our rationale, thus, is that neuron-specific modulations select a concatenation of decision boundary segments with constructible normal vectors that optimally approximates the desired decision boundary. By consequence, input weight vectors are preferentially constrained to the subspace of the data, so that all constructible normal vectors also lie within this subspace (Fig 4C). When there is no *a priori* information on the decision boundaries that might be drawn through the data, a reasonable heuristic for the constructible normal vectors is that they approximate the set of difference vectors between data samples. In turn, decision boundaries can be seen as a concatenation of segments with normal vectors that are close to difference vectors between nearby, but differently classified data samples (Fig 4D). Consequently, by aligning the set of constructible normal vectors of the network to the set of difference vectors between data samples, we ensure that constructible normal vectors lie within the data subspace, and that they constitute useful putative decision boundary directions.

**Figure 4.**
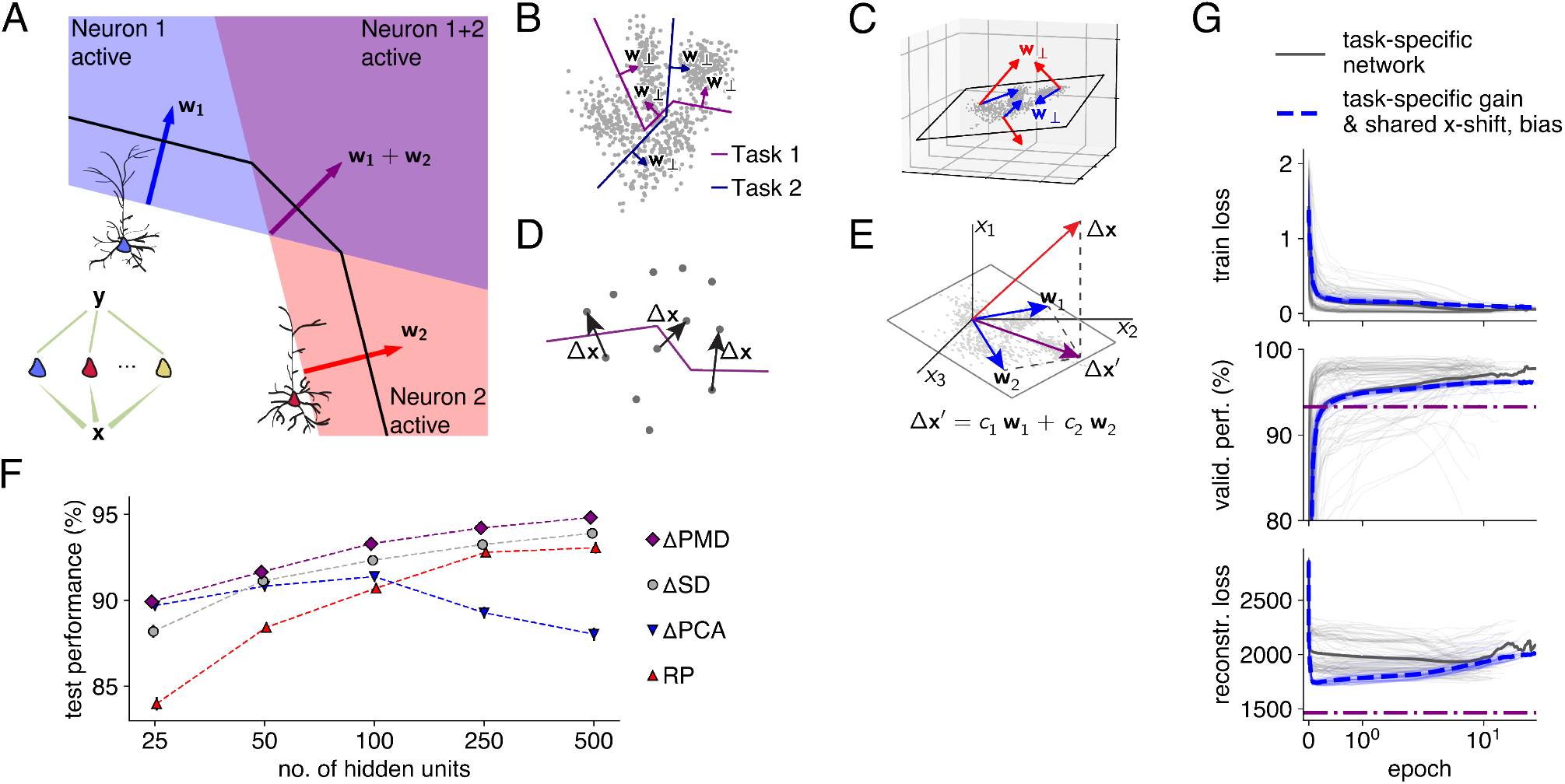
Properties of feedforward weights for networks to perform well in concert with neuron-specific modulations. **A:** The normal vectors associated with segments of the decision boundary capture the local features that the network uses to make decisions about data sample identities (note that with ReLU units, the decision boundary consists of linear sections). In any network architecture (here, a single hidden layer, inset), these normal vectors are weighted sums of the input weight vectors to the first layer neurons. **B:** To learn a multitude of tasks with the same feedforward weights, task-relevant normal vectors to the decision boundaries must be constructible with the network. **C:** Normal vectors to decision boundaries must be constrained to the subspace of the input data. Normal vectors outside this subspace have components orthogonal to it, which do not add useful directions for decision boundaries. **D:** Generic decision boundaries can be constructed by concatenating segments with normal vectors close to difference vectors between close, but differently classified data points. **E:** Combining considerations A-D, we investigate loss functions that minimize the difference between, on the one hand, difference vectors between data points and, on the other hand, their projections on the subspace spanned by the weight vectors to the first layer neurons. **F:** Performance on multitask EMNIST (averaged over five initialization seeds) as a function of layer size for networks with neuron-specific modulations and feedforward weights given by ΔPMD (purple diamonds), ΔSD (grey circles), ΔPCA (blue triangles), and random projections (RP, red circles). **G:** Train loss (top), validation performance (together with ΔPMD test performance as in F for comparison, middle) and reconstruction loss (min_*C*_ ∥Δ*X* −*CW*∥, bottom) during fully supervised learning (as in Fig 3) of the input weight matrix *W*, for an architecture with one hidden layer of 100 units, and for task-specific networks (grey) and neuron-specific modulations (blue, dashed). The reconstruction loss was evaluated on *W* during supervised training, and compared with the case were *W* was given by ΔPMD (purple, dash-dotted).

To achieve such alignment, we minimize the residual min_**c**_ ∥Δ**x** − **c***W*∥_2_ between any given difference Δ**x** and its optimal reconstruction as a linear combination of input weight vectors (the rows of the input weight matrix *W*, Fig 4E) with respect to *W* for a representative set of differences

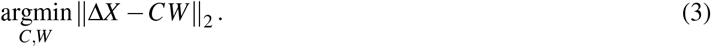

In this reconstruction loss, Δ*X* is a matrix with as rows the difference vectors and *C* the matrix with as rows the optimal coefficients **c**. We minimize (3) in three different ways (Table S1, Fig S2A). First, the optimum of (3) without regularizer or constraint (for lower hidden layer dimensionality *k* than input dimensionality *n*) is given by the principal components of Δ*X* (ΔPCA), and this problem can be solved in a biologically plausible manner through Hebbian learning rules^31^. Second, to encourage alignment between input weight vectors and difference vectors, we ask that any given Δ**x** can be expressed with few weight vectors. We achieve this by adding an L1-regularization term *λ* ∥*C*∥_1_ to (3). Thus (3) becomes the canonical sparse dictionary learning problem (ΔSD)^52,53^, which can also be solved by neural networks with biologically plausible Hebbian learning rules^32^. Finally, we encourage input weight vectors to capture local pixel correlations. We achieve this by placing an L1 constraint on ∥**w** _*j*:_ ∥ _1_ ≤ *ε* on the rows of *W*, next to an L1 constraint ∥**c**_: *j*_∥ _1_ ≤ *δ* for the columns of *C*. This doubly constrained minimization is known as the penalized matrix decomposition (ΔPMD)^54^.

We then embed the obtained feedforward weight matrices *W* in a network architecture with a single hidden layer of gain-modulated neurons, with shared x-shift and bias. The hidden neurons target a single gain-modulated output unit through identical feedforward weights. We find that solving (3) for differences between data samples instead of the data samples themselves generally results in a performance increase when combined with neuron-specific modulations to solve multitask EMNIST (Fig S2B). Assessing the relationship between input dimensionality (*n* = 784) and dimensionality of the hidden layer (number of hidden neurons *k*), we find that ΔPCA performs well for low numbers of hidden neurons, but that task performance saturates quickly and decreases for *k* ≥ 100 (Fig 4F, blue). This result is in agreement with our theoretical considerations: when the effective dimensionality of the data is reached, further orthogonal components do not contribute usefully to the decision boundary, as they lie outside of the subspace of the input data. In contrast, using random projections (RP) in *W* by sampling from a Gaussian distribution results in performances that increase strongly with *k* (Fig 4G, red). This can be understood by considering that with increasing numbers of random vectors, it becomes more likely that their linear combinations can approximate difference vectors between data points. Finally, we find that ΔPMD reached the highest performances for all *k* (Fig 4G, purple). These weight vectors being sparse likely facilitates learning performant sets of neuron-specific modulations, as up-or down-regulating a specific hidden neuron influences only a localized area of the input space. By consequence, neurons with receptive fields in other areas of the input space do not need re-adjustment, whereas neurons with non-local receptive fields would need to be re-adjusted. In these optimizations, the shared x-shift and bias, as well as the learning rate, were optimized through an evolutionary algorithm for each configuration separately (Fig S2C).

Finally, we investigate whether the supervised approach trained purely on the classification loss (as in Fig 3) also minimizes (3), to understand the extend to which supervised training adapts the input weight matrix to the subspace of the data, and how that depends on the task set. To that end, we compare the value of the residual min_*C*_ ∥Δ*X* −*CW*∥ of the reconstruction loss (3) during training of *W* in a fully supervised fashion (as in Fig 3), with the reconstruction loss when *W* was given by ΔPMD. We assess the case where the entire network was task-specific (feedforward weights included) and the gain-modulated case, and find that the sharp initial reduction in train loss (i.e. the classification loss) is associated with a sharp reduction in reconstruction loss (Fig 4H). The reconstruction loss for the gain-modulated network, where a single feedforward weight matrix has to contribute to solving a multitude of tasks, decreased to a value much closer to the ΔPMD optimum than the task-specific network average, where the feedforward weight matrix only has to solve a single task. Thus, supervised training forgets the initialization by adapting feedforward weights to the data subspace, and then fine tunes to the specific set of tasks, as validation performance during the later phase increases from the unsupervised value (obtained with ΔPMD) to its maximum. The unsupervised approach on the other hand is generalist, and admits any task that might be defined post hoc, at the cost of forgoing task-specific fine tuning.

### Spiking networks with biophysically realistic dendritic branches learn task switching online

To investigate whether our biologically inspired learning strategy holds with biophysically realistic neurons, we implement a spiking network of cortical pyramidal cell models (Fig 5A). We simplify the L5 PC model using the method developed in our previous work^55^, obtaining a model that was computationally sufficiently inexpensive to permit the network to be run over long timescales, thus allowing us to present a large amount of inputs. The output neuron is a single compartment model, obtained by only fitting the soma of the full L5 PC model, whereas the hidden layer consists of 100 neurons, each equipped with 20 dendritic compartments where context-modulating AMPA+NMDA synapses impinge (Fig 5A, blue). Feedforward weights to the hidden neurons are given by either ΔPMD, ΔSD, PCA or RP, whereas all feedforward weights to the output neuron have the same value.

**Figure 5.**
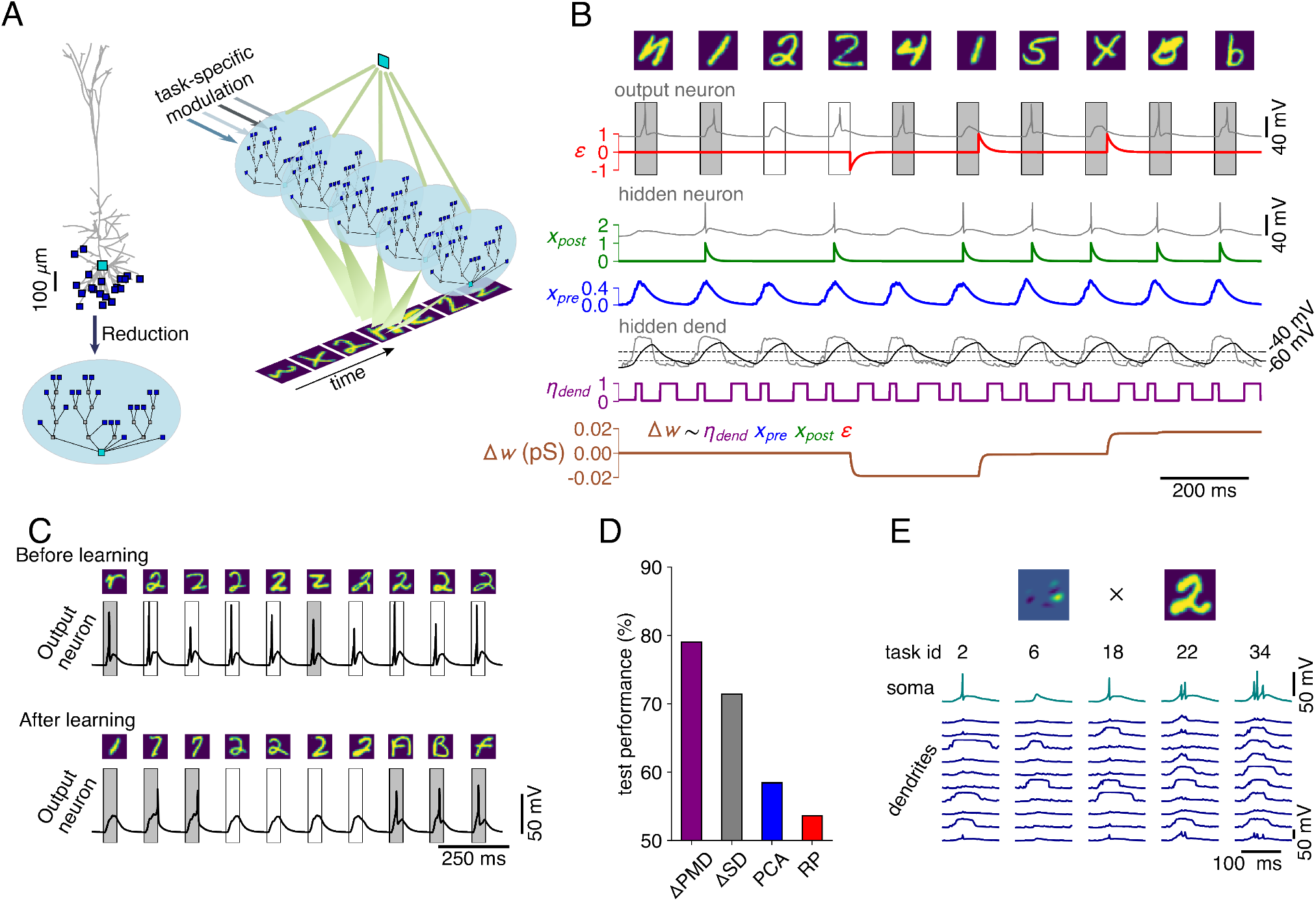
Dendritic branches learn to solve multitask EMNIST through a biologically plausible learning rule. **A:** To simulate a feedforward network consisting of neurons with biophysically realistic dendritic subunits for a sufficiently long time, we reduce the L5 PC and synapse configuration shown in Fig 1F^55^. We then connect these neurons to an output neuron – implemented as a single-compartment reduction of the same model – that learns to spike in response to a random sample and remain silent in response to a sample from the class to be recognised. **B:** Weight changes of dendritic synapses (brown, bottom) are computed as the product of a global error signal (red), a low-pass filter of the post-synaptic spikes (green), a low-pass filter of the pre-synaptic spikes (blue) and a voltage-dependent learning rate modulation (purple). **C:** Voltage trace of the output neuron before learning (top) and after learning (bottom), for the network with the ΔPMD feedforward weight matrix, during an examplar one-vs-all task (not spiking in response to ‘two’). **D:** Performance on multitask EMNIST of the resulting model for the different feedforward weight matrices, labels as in Fig 4F,G. **E:** Somatic voltage (teal) and a subset of dendritic voltages (blue) in a representative hidden neuron, for the same feedforward input (i.e the inner product of the input weight vector [top, left] and a randomly chosen data sample [top, right]) and five example tasks (left to right).

Learning at the dendritic synapses was orchestrated by an online error-weighted Hebbian plasticity rule during a continuous stream of inputs (Fig 5B). Because of our choice of architecture, with identical weights from all hidden neurons to the output neuron, this learning rule approximately follows the error gradient of the classification loss (methods). For each data sample, the pixel intensities are converted into short, Gaussian bursts of spikes (width of 6 ms), with spike numbers proportional to pixel intensity. These spikes are fed into feedforward synapses whose weights were scaled according to the matrices computed in the previous section. Conversely, the task context is encoded by a wide Gaussian burst (width of 20 ms), consisting of on average 60 spikes if the context is active and zero spikes otherwise. The first of the feedforward spikes opens a 50 ms window in which the output neuron should either generate an output spike – in response to a random sample – or generate *no* output spike – in response to a sample from the class to be recognized. In case of erroneous firing, a global error signal (Fig 5B, red) is relayed to the dendritic synapses of the hidden neurons. This error signal is then multiplied by a low-pass filter of the somatic spike output (Fig 5B, green), a low-pass filter of the presynaptic spike input (Fig 5B, blue), and a learning rate modulation (Fig 5B, purple) based on a low-pass filter of the local dendritic voltage (Fig 5B, black).

This network architecture solves multitask EMNIST, and tasks that are demonstrably not linearly separable, such as XOR (Fig S3). Initially, the output neuron fires indiscriminately, but learns to spike correctly during the target intervals (Fig 5C, Fig S3C,D, shaded boxes). Assessing network performances averaged over all 47 tasks (Fig 5D), we find that performance differences observed between alternative feedforward matrices for the artificial network architecture (Fig 4F,G) are exacerbated, with RP performing barely better than chance level. Thus, in the noisy and imprecise spiking system, it is all the more important that the feedforward weight matrix consists of localized receptive fields, well-adapted to the input data. Our ΔPMD matrix achieves this for multitask EMNIST. Finally, we assess the somatic and dendritic activity after learning in the same hidden neuron, for the same feedforward input, across different tasks (Fig 5E, in the ΔPMD-network). We find that between zero and three output spikes are emitted, depending on the precise dendritic state. Thus, this network successfully learns multitask EMNIST by expressing a different dendritic state for each task. These learned dendritic states modulate rapid feedforward processing to solve a multitude of tasks, supporting our central hypothesis.

### Task-modulated contrastive learning for stacking processing layers

Sensory processing in the brain is thought to proceed in a hierarchical manner through a number of processing layers^9,10^. Deep artificial networks also implement hierarchical processing through a stack of layers, the learning of which is orchestrated by error backprogagation^56,57^. Nevertheless, the question of whether this algorithm could plausibly be implemented in the brain is still a matter of debate^58^, in contrast to representation learning approaches such as PCA^31^ or SD^32^, which have biologically plausible implementations. These representation learning approaches, however, do not extract higher-order features when stacked in a deep network^51^. Furthermore, by introducing neuron-specific modulations to the hidden processing layers, the representation learning problem becomes even more complex, as now the hidden representations depend on task modulation.

Here, we propose a representation learning algorithm that does not rely on error backpropagation between layers, and where the task dependence of the hidden representations is an integral feature that improves generalization. As we have shown above, sparse feedforward connectivity is beneficial in concert with neuron-specific modulations. We therefore apply our algorithm to a convolutional architecture (Fig 6A), which by design features localized receptive fields adapted for visual processing^59^. Our representation learning approach takes inspiration from a successful contrastive learning (CL) algorithm^60^. In this algorithm, augmentations (e.g. occlusions, rotations, scalings and combinations thereof) are applied to the input data and the convolutional feedforward network creates hidden representations thereof. A multi-layer perceptron (CL-MLP) – applied to these hidden representations – is trained in concert with the convolutional feedforward weights to maximize similarity between representations if they originate from augmentations of the same input sample; conversely to maximize contrast if they originate from different input samples. In the original formulation^60^, the CL-MLP is applied once at the end of the feedforward pathway, and weight changes are orchestrated across layers by error backpropagation of the CL loss. Here, we construct our networks layer by layer by applying this algorithm in a layer-wise fashion. Hence, a local CL-MLP minimizes the CL loss to learn the feedforward weights between the previous and the current layer, and no error gradients propagate across layers (Fig 6A, see methods for details). After this CL phase, we learn task-specific gains for the hidden neurons in the current layer through a task-independent output unit (OU) that maximises classification performance in a supervised manner (Fig 6A, blue). To train feedforward weights to the next layer in the next CL phase, the task-modulations learned in the previous layers are treated as additional data augmentations, across which similarity has to be maximized (Fig 6B).

**Figure 6.**
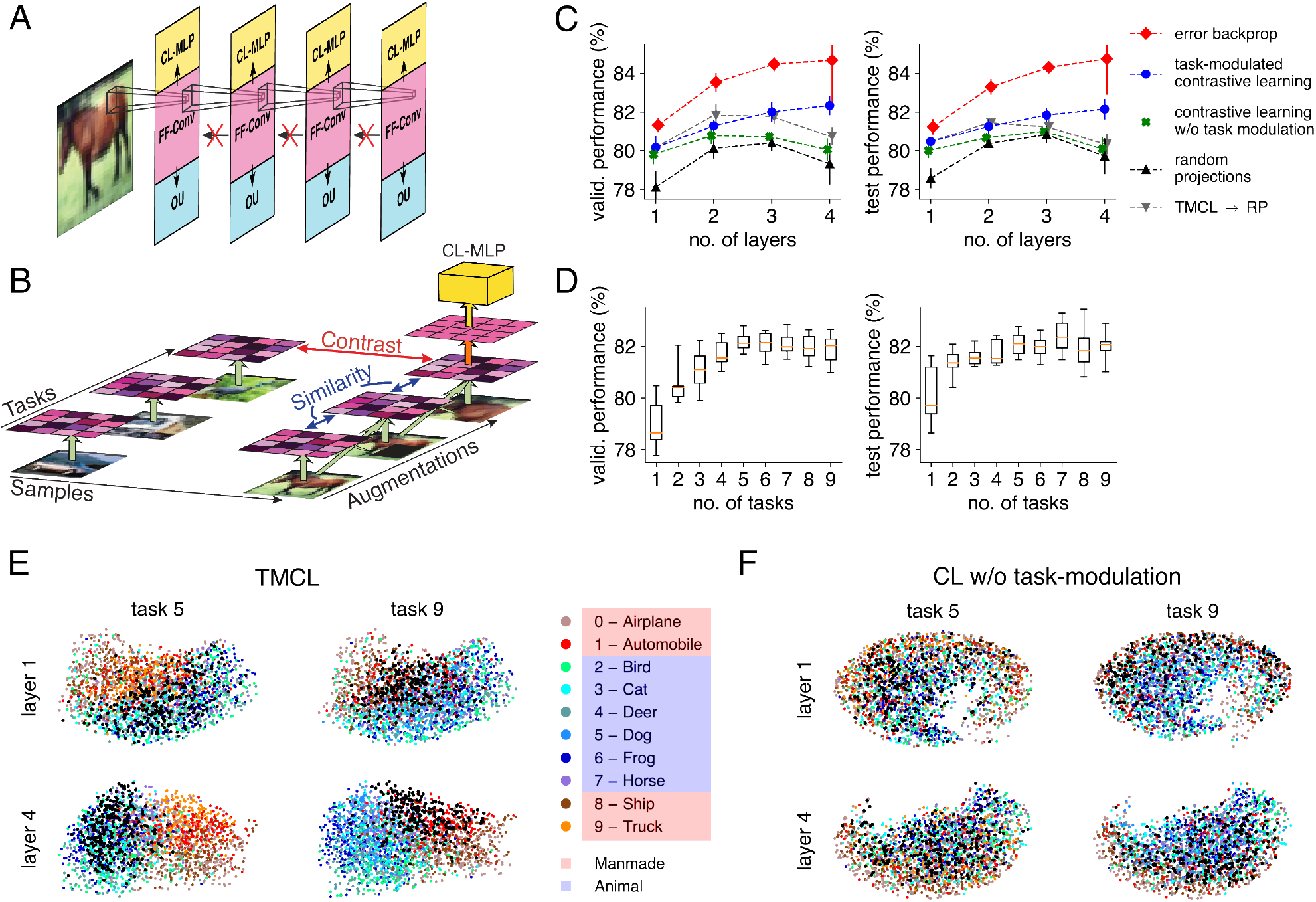
Hierarchical stacking of task-modulated convolutional layers. **A:** We train a stack of gain-modulated convolutional layers on multitask CIFAR-10 using a contrastive learning (CL) objective. Each layer consists of a CL multi-layer perceptron (CL-MLP, yellow) to implement the CL objective, a set of convolutional feedforward weights (purple), and an output unit (blue) to learn the task-specific gains. In this task-modulated contrastive learning (TMCL) paradigm, no error gradients flow back between layers. **B:** To learn the convolutional feedforward weights to the next layer (orange arrow), the CL-MLP maximizes contrast between representations in the last learnt layer that originate from different data samples, and similarity between representation that originate from augmentations (occlusions, scalings, rotations and combinations thereof) of the same data sample to which, additionally, different task-modulations are applied (green arrows represent the feedforward pathway up until the last learnt layer). **C:** Validation (left) and test (right) performances on multitask CIFAR-10 (averaged over five initialisation seeds) for the gain-modulated networks (with shared x-shift), with filters trained by: error backpropagation (red), TMCL (blue), contrastive learning without similarity prediction across task-modulations (green), given by random projections (RP, black), or RP stacked on top of a TMCL layer (grey). **D:** Validation (left) and test (right) performances of TMCL for networks with four layers, where during the TMCL phase the augmentations where fed to a subset of the task-modulations (ten random but distinct subsets where evaluated for each number of tasks). Median performance, orange; box denotes [Q1, Q3] over the ten subsets, whiskers [min, max]. **E:** UMAP projections of the hidden, task-modulated representations for the TMCL-trained network. Color code as in the legend, except that the class to be recognized is black (‘dog’ for task 5 and ‘truck’ for task 9). **F:** UMAP projections of the hidden, task-modulated representations for the TMCL without task-modulation. Color code as in E.

We test our task-modulated contrastive learning (TMCL) algorithm on multitask CIFAR-10^61^, and find that network performance, averaged over all tasks, increases with the number of layers (Fig 6C, blue). To establish a performance envelope, we test equivalent network architectures trained in a fully supervised manner through end-to-end error backpropagation (Fig 6C, red). The performance of networks with RP starts at lower values, and does not increase as much, or even decreases, across layers (Fig 6C, black). Similarly, stacking RP layers on top of a TMCL layer does not increase performance across multiple layers (Fig 6C, grey). Removing similarity maximization across task representations from TMCL also abolishes the performance increase with stacking (Fig 6C, green). We furthermore assess network performance while using different numbers of tasks during the TMCL phase. For each number of tasks, we select ten random but distinct subsets containing that amount of tasks, and use only those tasks as hidden augmentations. We then evaluate network performance across all tasks, and find that performance increases with subset size (Fig 6D), indicating that using representations with many tasks improves generalization.

Finally, we visualize how the network constructs and modulates hidden representations, to investigate whether high-level information is extracted across layers. We apply the uniform manifold approximation and projection (UMAP^62^), a non-linear visualisation method, to TMCL-generated hidden representations. In the first layer, at most a general distinction between manmade objects (red shades) and animals (blue shades) can be observed, while in the fourth layer individual classes appear in a localized pattern (Fig 6F). Such localized patterns cannot be distinguished for CL without task-modulations (Fig 6F) or for RP (Fig S4).

## Discussion

In this work, we have laid out a biophysical theory of how contextual modulations can be achieved in the brain. We have shown that the dendritic trees of neurons are ideally suited to integrate contextual inputs, resulting in concerted gain and threshold modulations to the IO relationship of individual neurons. We have then demonstrated that such task-modulations achieve performant transfer learning. Furthermore, the architecture of our models, with task-modulations that are applied prior to the non-linear activation – i.e. the spiking output in networks of biophysically realistic neurons – and identical feedforward weights to the output neuron, allows a Hebbian plasticity rule modulated by a global error signal to learn non-linear task-specific decision boundaries. Finally, we have shown that task-modulations to hidden layers augment sensory representations, facilitating the extraction of high-level features through contrastive learning.

While the component of our task-modulated contrastive learning approach that learns task-modulations can be implemented in a biologically plausible fashion, as shown through our network model with realistic dendritic subunits, the contrastive learning step in our study relies on precise error backpropagation through the CL-MLP. However, a contrastive learning algorithm has recently been proposed in the context of predictive coding that relies solely on Hebbian learning rules^33^. This algorithm shows that contrastive learning could be implemented in a self-supervised manner, by neurons connecting locally to principal feedforward cells (L5 PCs), and using gaze information to assess whether similarity or contrast have to be maximised.

A puzzling observation, first discovered in high-level areas^63,64^ and later also in early sensory regions^65–67^, is that the participation of a neuron in the representation of a sensory stimulus changes over time. This representational drift raises questions about the framework of classical representation learning, and about how stable perception can be achieved^68,69^. As we show, changes to the sensory representation could help in extracting high level information in further processing layers. The drift itself could be a manifestation of changes in the internal mental state – encoded on dendritic trees and thus invisible in most imaging experiments.

Feedforward processing needs to be rapid, for instance to initiate evasive action when a threat is identified, while contextual modulation likely proceeds on a slower time scale, for instance to bias the feedforward pathway towards detection of relevant threats given an environment. We have linked this difference in time scales to the underlying biophysical processes: the short duration of somatic spikes (1-5 ms) and by extension the whole feedforward pathway (100-150 ms)^34,35^ in comparison to the duration of dendritic spikes (50-100 ms, or possibly longer^42,44^). These temporal scales match the frequency bands associated with feedforward processing (gamma, 60-80 Hz) and top-down processing (alpha-beta, 10-20 Hz) observed across a large range of tasks and stimuli^70–75^.

In the brain, the contextual signal to a neuron is likely a rich combination of cross-modal information, recurrent information about the recent past, and top-down signals about high-level goals, behavioral state and environment characteristics. That these signals impinge on dendritic branches, which function as semi-independent feature detectors^76^, has the added benefit of suppressing noise by preventing spurious activations by random subsets of dendritic inputs^77^. Furthermore, distinct contextual pathways may target distinct loci on the dendritic tree. Local recurrent connections, e.g. L5 PC to L5 PC or L5 PC to L2/3 PC to L5 PC^78^, target basal and proximal apical dendrites^79^, and may relay information about the recent past as a context for the present. Axons carrying top-down signals target L5 and L6^40^, where they generate NMDA spikes in basal dendrites of L5 PCs, as well as L1^80^, where they generate Ca^2+^ spikes in the apical dendrites of L5 PCs^42,81^. Whereas we have focused on NMDA-spikes in this work, Ca^2+^ also provide gain modulation over long time-scales and are thus suited for implementing contextual adaptation^82^. Note furthermore that top-down signals targetting L1 need not remain constrained to the apical dendrites of L5 PCs. These signals also target GABAergic somatostatin-expressing (SST) and vasoactive intestinal peptide-expressing (VIP) interneurons^80^, which in turn provide inhibition and disinhibition, respectively, to the basal dendrites of L5 PCs^36^, and powerfully regulate NMDA-spike genesis and duration^83,84^.

Taken together, our work reframes feedforward processing in the brain as a fundamentally adaptable process, steered dynamically by contextual inputs that modify the dendritic state. Our theory matches environmental constraints to the underlying biophysical layout, and may help to explain diverse observations, such as the frequency bands associated with feedforward and top-down processing, and the apparent instability of sensory representations.

## Acknowledgements

We thank Angela Fischer and Dr. Sanne Rutten for generous help with the schematic diagrams in this paper. This work was supported by the Helmholtz Portfolio theme Supercomputing and Modeling for the Human Brain, the Excellence Strategy of the Federal Government and the Länder [G:(DE-82)EXS-PF-JARA-SDS005, G:(DE-82)EXS-SF-neuroIC002] and by the Swiss National Science Foundation (Grant 180316 to WS). The project received additional funding from the European Union’s Horizon 2020 research and innovation programme under Specific Grant Agreement Nos. 720270, 785907 and 945539 (Human Brain Project SGA1-3). The authors gratefully acknowledge the computing time granted by the JARA Vergabegremium and provided on the JARA Partition part of the supercomputer JURECA at Forschungszentrum Jülich (Grant 25241 to WW). We also thank the Insel Data Science Center for using their HPC Cluster.

## Methods

### Biophysical modelling

The morphology, ion channels and physiological parameters for the L5 PC model were taken from Hay et al.^45^ and implemented in the NEURON simulator^85^. Contextual and background synapses were conductance based, either containing AMPA+NMDA (excitatory) or GABA (inhibitory) receptors. AMPA and GABA receptors were implemented as the product of a double exponential conductance profile^86^ *g* with a driving force:

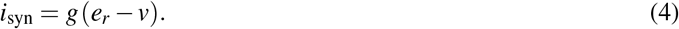

AMPA rise resp. decay times were *τ*_*r*_ = 0.2 ms, *τ*_*d*_ = 3 ms and AMPA reversal potential was *e* = 0 mV. For GABA, we set *τ*_*r*_ = 0.2 ms, *τ*_*d*_ = 10 ms and *e* = −80 mV. N-methyl-D-aspartate (NMDA) currents^87^ were implemented as:

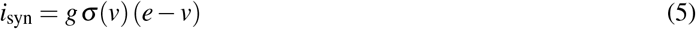

with rise resp. decay time *τ*_*r*_ = 0.2 ms, *τ*_*d*_ = 43 ms, and *e* = 0 mV, while *σ* (*v*) – the channel’s magnesium block – had the form^88^:

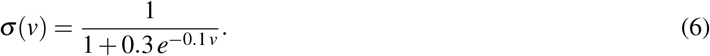

The weight of a synapse signifies the maximum value of its conductance profile. For an AMPA+NMDA synapse, the weight is the maximal value of the AMPA conductance profile, and the maximal value of the NMDA profice is twice that of the AMPA window (NMDA ratio of 2). Feedforward synapses where current based, with a dual exponential current profile with *τ*_*r*_ = 0.2 ms and *τ*_*d*_ = 3 ms. The excitatory synaptic weight corresponded to the maximal value of the current profile. For inhibitory feedforward synapses, the weight was negative and corresponded to the minimum value.

For dendritic modulation, we identified 20 dendritic compartments suited for semi-independent NMDA-spike generation (e.g. avoiding compartments on sister branches, Fig 1F) and equipped each of these compartments with an AMPA+NMDA synapse. Both the feedforward and shunt inputs targeted the somatic compartment. The 20 dendritic compartments as well as the soma were also equipped with an AMPA and GABA background synapse. All parameters for the simulations are shown in Table 1.

**Table 1.** Simulation-specific parameters to investigate somatic and dendritic modulation (Fig 1,2). Note that for dendritic modulation, we set the number of compartments with NMDA-spikes *n*_*c*_ by targetting *n*_*c*_ compartments with an AMPA+NMDA synapse of high weight and 20 −*n*_*c*_ compartments with an AMPA+NMDA synapse of low weight, and delivered the same amount of input spikes in each case. For somatic modulation, the weight of the shunt synapses remained the same, but the number of input spikes was modified. Note that the burst width for somatic modulation was enlarged to account for the shorter time-scale of the GABA conductance window compared to the NMDA conductance window.

|  | **Dendritic modulation** | **Somatic modulation** |
| --- | --- | --- |
| Weight exc (inh) feedforward synapses | 0.06 (-0.06) nA | 0.06 (-0.06) nA |
| Feedforward inputs burst width | 6 ms | 6 ms |
| No. of AMPA+NMDA inputs | 60 |  |
| High AMPA+NMDA weight | 0.08 nS |  |
| Low AMPA+NMDA weight | 0.001 nS |  |
| No. of comps with NMDA spike | {0, 3, 6, 9, 11, 14, 17, 20} |  |
| Shunt weight |  | 5 nS |
| No. of shunt inputs |  | {0, 20, 40, 60, 80, 100, 120} |
| Contextual inputs burst width | 20 ms | 40 ms |
| Soma comp AMPA (GABA) background rate | 100 (200) Hz | 100 (200) Hz |
| Dend comps AMPA (GABA) background rate | 60 (120) Hz | 60 (120) Hz |
| Weight AMPA (GABA) background synapses | 1 (1) nS | 1 (1) nS |

To measure the membrane conductance change induced by the contextual inputs (Fig 1H), we follow Haider et al.^46^ and measure the time-dependent electrode current in voltage clamp (*i*(*t*; *v*_*h*_), with *v*_*h*_ the holding potential) for two different holding potentials: *v*_*h*1_ = −80 mV and *v*_*h*2_ = 0 mV. Following the Ohmic relationship *g*(*t*)(*v*_*h*_ − *v*_0_) = *i*_*h*_(*t*; *v*_*h*_), with *v*_0_ the equilibrium potential, we find the membrane conductance as

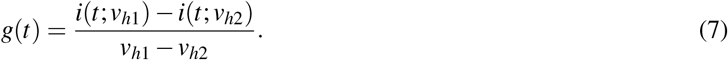

To visualize *g*(*t*) as in Fig 1H, we subtract the pre-stimulus baseline and plot the min-max envelope obtained over 10 trial runs.

### IO curve parameter fitting

To obtain the IO curves, we change the number of feedforward inputs in each burst (weights as in Table 1), and either have only excitatory inputs (no. of feedforward inputs *>* 0) or inhibitory inputs (no. of feedforward inputs *<* 0). For each number of inputs, we present ten independently sampled bursts featuring that number of inputs, and measure the average number of output spikes generated in response.

To fit the IO curves (Fig 2), we a use linear least squares fit (argmin_**u**_ ∥*A* **u** − **y**∥_2_) both for the shared gain and shared bias cases, but construct the feature matrix *A* and parameter vector **u** differently. On the domain where *y >* 0, the IO curve is

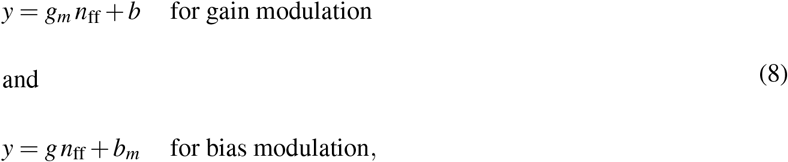

with *n*_ff_ the number of feedforward inputs. For M modulation levels, we have parameter vectors **u** = (*g*_1_, *g*_2_, …, *g*_*M*_, *b*) for gain modulation and **u** = (*g, b*_1_, *b*_2_, …, *b*_*M*_) for bias modulation, and corresponding feature matrices

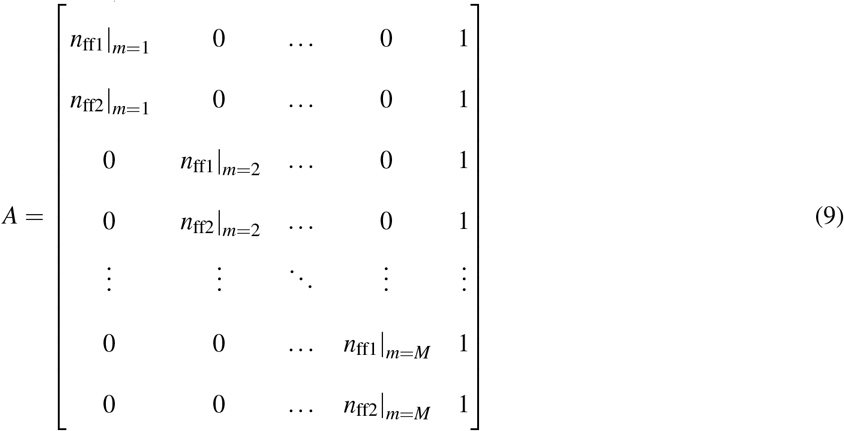

for gain modulation and

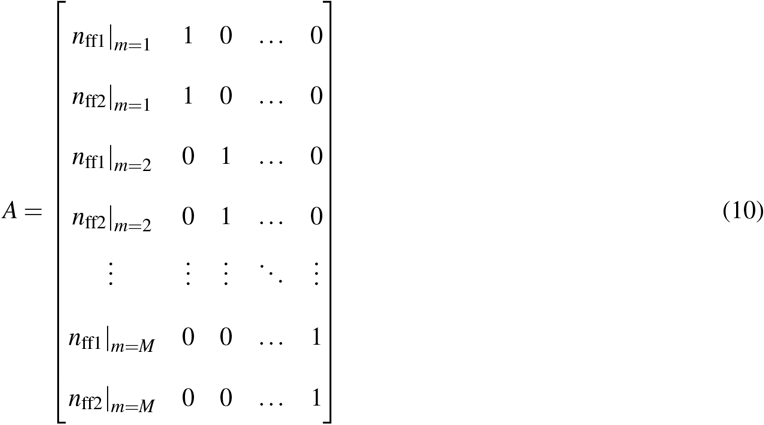

for bias modulation, with *n*_ff1_|_*m*_ corresponding to the number of feedforward inputs where the output transitioned from 0 to 1 spikes and *n*_ff2_|_*m*_ from 1 to 2 spikes, given a modulation level *m*. The output vector **y** = (0.5, 1.5, 0.5, 1.5,…, 0.5, 1.5) was identical in both cases. To extend gain modulation with a shared x-shift, we modified the entries in the gain modulation feature matrix *A* from *n*_ff*i*_|_*m*_ to *n*_ff*i*_|_*m*_ − *x*_shift_ (*i* = 1, 2, *m* = 1, …, *M*) and minimized the fit residual min_**u**_ ∥*A*(*x*_shift_) **u** − **y**∥_2_ over the *x*_shift_ values.

### Supervised learning in the fully connected networks

The fully connected network architectures with neuron-specific modulations consisted of a fixed number of layers of equal size, the final layer connected to a single output unit. The neuron-specific modulations were implemented in an abstract fashion, as task-specific parameters, although it is straightforward to extend the network architecture to implement the modulations as synaptic inputs. The networks were trained through end-to-end error backpropagation. For the networks with task-specific outputs, the hidden units had unit gain (fixed) and a trained bias shared across tasks, and as many output units as there were tasks. The hidden layer neurons had ReLU transfer functions and the output unit a tanh(*·*) transfer function. The networks were implemented in PyTorch^89^, trained on batches of size 1000 using Adam^90^ and we computed the mean squared error between target and network output as loss. The targets were defined as either −1 or +1, and a custom sampler assured batches consisted of an equal number of samples from each task and task-class. The training was interrupted with an early stopping criterion with a patience of 5 epochs on the validation performance evaluated after each epoch on a validation set of 47932 samples, or after 50 epochs, whichever came first. Independent learning rates for shared and task-specific parameters were optimized for each form of multitask leaning separately with an iterative grid search (Fig S1C). Performances are measured by averaging over all tasks, and by additionally averaging over twenty initialization seeds (error bars show standard deviation of task-performance across seeds, averaged over all tasks).

### Decision boundary normal vectors

To explain our approach, we consider the pre-activation *a* : ℝ^*n*^ → *ℝ* : **x** → *a*(**x**) of a neuron in a feedforward network as a function of the sensory input **x** (the activation *y* being given by *y*(**x**) = *σ* (*a*(**x**))). Biophysically, this quantity most closely corresponds with the somatic voltage under Na^+^-channel blockage; it is the aggregate of all inputs and when it crosses a threshold, the neuron would emit a spike if the non-linear activation function (the Na^+^-channels) were applied. The neuron may be active (*a*(**x**) *>* 0) or inactive (*a*(**x**) *<* 0), and its decision boundary on the input domain is given by the set D = {**x**_D_ ∈ *ℝ*^*n*^ | *a*(**x**_D_) = 0}. In a small enough region around a point **x**_D_ ∈ D, *a*(**x**) can be approximated as being linear:

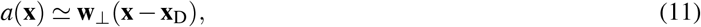

and 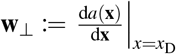 is the local normal vector of the decision boundary. This normal vector is always a linear sum of the input weight vectors **w** _*j*_ to the first layer neurons (*j* = 1, …, *k*, with *k* the number of neurons in the first layer), and is perpendicular to the local decision boundary (see methods), thus capturing the local input features that *a* uses to make a decision about whether to become active close to **x**_D_ (Fig 4A).

First, we note that for input **x** ∈ D on the decision boundary D close enough to **x**_D_, *a*(*x*) = 0, and thus, following (11):

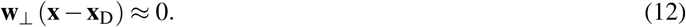

By consequence, **w**_⊥_ is perpendicular to the local decision boundary and indeed its normal vector.

Second, we compute **w**_⊥_ explicitly, and to that purpose write the preactivation *a* of a neuron in layer K of the network as:

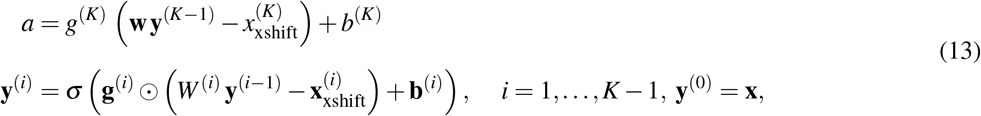

with **y**^(*i*)^ the neural activations in layer *i, σ* : ℝ → *ℝ* the neural activation function applied element-wise to its inputs, *W* ^(*i*)^ the weight matrix from layer *i* − 1 to layer *i*, **g**^(*i*)^, 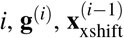. **b**^(*i*)^ the gains, x-shifts resp. biases in layer *i, w* the weight vector to the neuron and *g, x*_xshift_ resp. *b* its gain, x-shift and bias. **w**_⊥_ is then found as

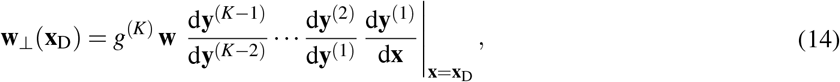

with d**y**^(*i*)^*/*d**y**^(*i*−1)^ the Jacobian matrix of **y**^(*i*)^ with respect to **y**^(*i*−1)^. This Jacobian is given by

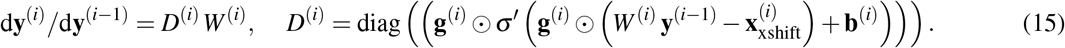

Thus, we find for (14):

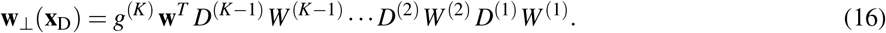

By rearranging this matrix product, we obtain

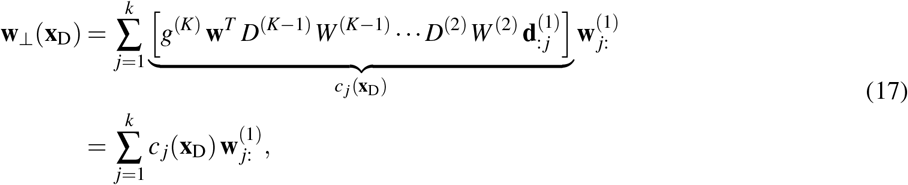

a linear weighted sum of the inputs weight vectors (with 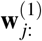 the *j*’th row of *W* ^(1)^ and 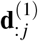 the *j*’th column of *D*^(1)^).

### Unsupervised learning of weights combined with supervised gain modulation

The unsupervised optimization problems for PCA, ΔPCA, SC, SD, and ΔSD are solved using Scikit-learn^91^, and for PMD and ΔPMD we use a custom implementation. The networks consisted of a single hidden layer, with weights resulting from optimising the loss functions in Table 2 (for RP, weights were drawn from a Gaussian distribution and then normalized so that 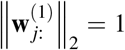. Weights to the output unit were uniform, and also normalized to have unit Euclidean norm 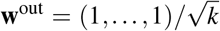, with *k* the dimensionality of the hidden layer). Activation functions, target outputs, the optimizer and the way in which performance was measured were identical to the fully supervised networks.

**Table 2.**
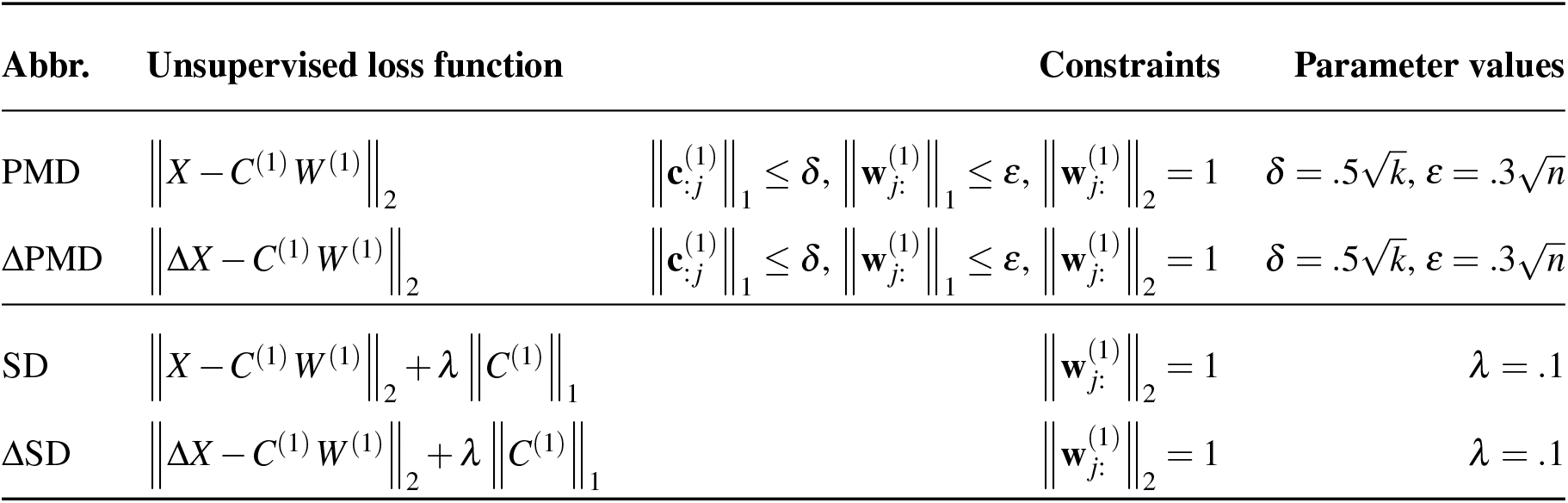
The unsupervised loss functions with their constraints and parameter values. *C*^(1)^ is commonly referred to as the sparse code (SC) and *W* ^(1)^ as the sparse dictionary (SD). Note that Δ*X, X* ∈ *ℝ* ^*k×n*^.

| Abbr. | Unsupervised loss function | Constraints | Parameter values |
| --- | --- | --- | --- |
| PMD | $\ X - C^{(1)} W^{(1)}\ _2$ | $\ \mathbf{c}_{:j}^{(1)}\ _1 \leq \delta, \ \mathbf{w}_{j:}^{(1)}\ _1 \leq \varepsilon, \ \mathbf{w}_{j:}^{(1)}\ _2 = 1$ | $\delta = .5\sqrt{k}, \varepsilon = .3\sqrt{n}$ |
| ΔPMD | $\ \Delta X - C^{(1)} W^{(1)}\ _2$ | $\ \mathbf{c}_{:j}^{(1)}\ _1 \leq \delta, \ \mathbf{w}_{j:}^{(1)}\ _1 \leq \varepsilon, \ \mathbf{w}_{j:}^{(1)}\ _2 = 1$ | $\delta = .5\sqrt{k}, \varepsilon = .3\sqrt{n}$ |
| SD | $\ X - C^{(1)} W^{(1)}\ _2 + \lambda \ C^{(1)}\ _1$ | $\ \mathbf{w}_{j:}^{(1)}\ _2 = 1$ | $\lambda = .1$ |
| ΔSD | $\ \Delta X - C^{(1)} W^{(1)}\ _2 + \lambda \ C^{(1)}\ _1$ | $\ \mathbf{w}_{j:}^{(1)}\ _2 = 1$ | $\lambda = .1$ |

We optimized gains for each task separately by performing gradient descent with batches of 100 samples, and performed an evolutionary meta-parameter optimization using DEAP^92^ to find optimal values for x-shift, bias, and learning rate by maximizing validation performance on a subset of 10 tasks (Fig S4C).

To construct Fig 4G, we evaluate the residual min_*C*_ ∥Δ*X* −*CW*∥ of the reconstruction loss (3) during supervised training without regularizer or constraint (i.e. the least mean squares optimum with respect to *C*) on matrices Δ*X* containing 1000 differences.

### Loss gradient with respect to task gains as a Hebbian learning rule with global error modulation

Here, we show that in the network architecture described in the previous section, the error gradient with respect to the task-dependent gains can be expressed as a Hebbian learning rule modulated by a global error signal. We interpret task-dependent gains to the hidden neurons with activity *y*_*i*_ (*i* = 1, …, *k*) as synaptic inputs originating from task-encoding neurons. With *g*_*i,t*_ the weight of the connection to neuron *i* associated with task *t*, and *z*_*t*_ ∈ {0, 1} the activity of the associated task-encoding neuron (1 if the task is active and 0 otherwise), the total gain is *g* = ∑_*t*_ *g*_*i,t*_ *z*_*t*_.

The structure of these networks then becomes

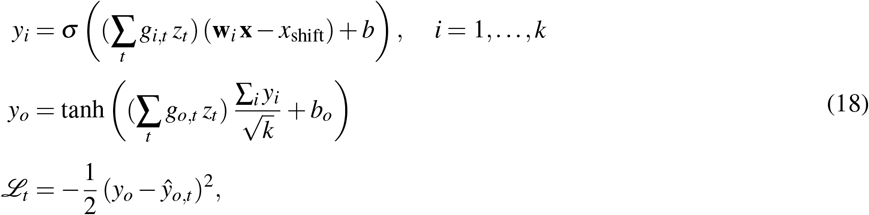

with *y*_*o*_ the activity of the output neuron, *b*_*o*_ its bias and *g*_*o,t*_ the weight of the task-gain connection to the output neuron. *ŷ*_*o,t*_ ∈ −1, 1 is the task-dependent target value for a given input sample. Computing the gradient of the task-loss ℒ _*t*_ with respect to the task-gains in the hidden layer yields

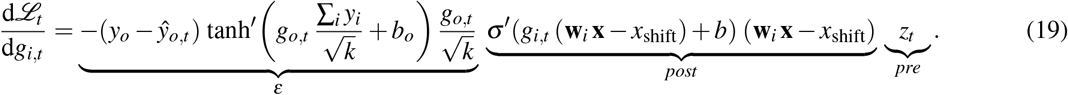

One crucial observation here is that since the feedforward weights to the output unit are all identical, they do not modify the error signal in a neuron-specific manner. By consequence, the error signal to the hidden layer is global, i.e. identical for all hidden neurons. A second crucial observation is that while it may seem as if the *post* factor could change the sign of the gradient, if

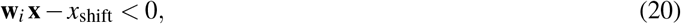

this is actually impossible with the ReLU activation function (which results in *σ′* ∈ {0, 1}), and positive gains together with negative bias, as obtained from the L5 PC model. Indeed, for the neuron to be active (*σ′* = 1), the inputs need to be sufficiently strong, so that

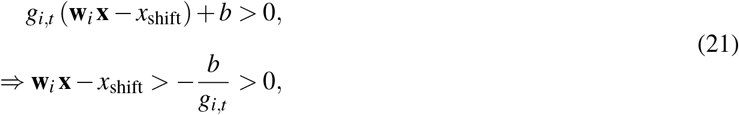

where the second inequality holds because we have negative bias and positive gains. Thus, whenever (20) is satisfied, the neuron is inactive (*σ′* = 0) and **w**_*i*_ **x** − *x*_shift_ does not influence the update of *g*_*i,t*_. We will leverage this fact in the spiking neural network, where we replace the *post* factor by a low pass filter of the somatic output spikes to obtain a learning rule that follows the approximate gradient.

### Spiking network

In order to simulate the network model, which consisted of one hidden layer with 100 neurons, over sufficiently long time scales to allow significant learning, we reduced the L5 PC model by retaining only the dendritic compartments and the soma (Fig 5A), using the method proposed by Wybo et al.^55^ to conserve dendro-somatic response properties. We then equip each dendritic compartment with an AMPA+NMDA synapse for each context – 47 in multitask EMNIST (Fig 5) and 14 for the boolean tasks (Fig S3) – whose weight evolved according to

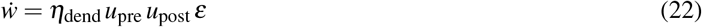

with *η*_dend_ a learning rate dependent on a low-pass filter *u*_dend_ of the local dendritic voltage *v*_dend_, *u*_pre_ a low pass filter of the input spikes to the contextual synapse, *u*_post_ a low pass filter of the output spikes and *ε* a global error signal, implemented as the low pass filter of an error pulse whose amplitude *a*_*e*_ was proportional to the difference between the number of generated output spikes in the ‘reward window’ and the expected number (i.e. 0 or 1). This 50 ms reward window opened upon the arrival of the first feedforward input spike associated with a data sample. A delta pulse with amplitude *a*_*ε*_ was then injected into *ε* at the closure time *t*_*e*_ of this reward window. Summarizing the above, we had the following:

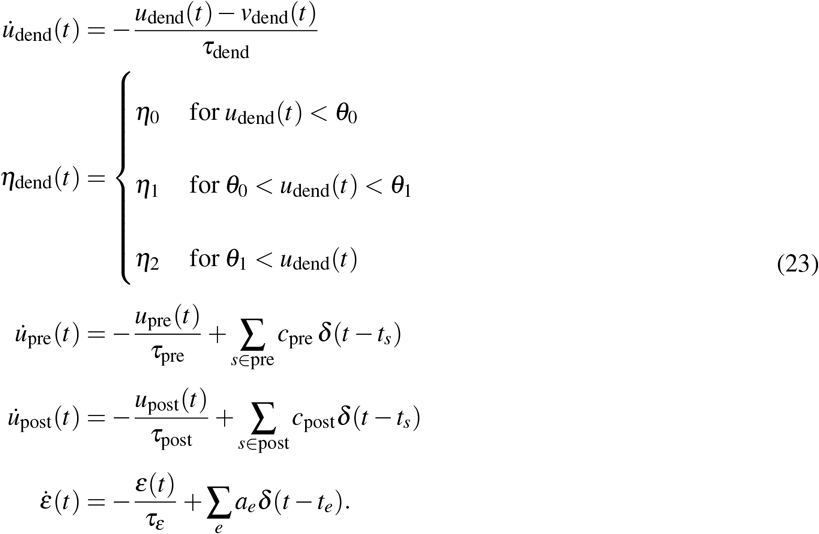

Note that *η*_dend_, *u*_dend_, *u*_pre_ and *u*_post_ are all specific to the synapse, whereas *ε* is global and shared across all synapses. Model parameters are summarized in Table 3.

**Table 3.** Parameters used in the network model with biophysically realistic neurons.

| Plasticity rule parameter | Parameter value |
| --- | --- |
| $\tau_{\text{dend}}$ | 25 ms |
| $\eta_0, \eta_1, \eta_2$ | $10^{-5}$ nS/ms, $10^{-4}$ nS/ms, $5 \times 10^{-6}$ nS/ms |
| $\theta_0, \theta_1$ | -50 mV, -10 mV |
| $\tau_{\text{pre}}, \tau_{\text{post}}$ | 25 ms, 10 ms |
| $c_{\text{pre}}, c_{\text{post}}$ | 1/60, 1 |
| $\tau_{\epsilon}$ | 15 ms |
| Network parameter | Parameter value |
| No. of feedforward inputs per burst | 0-10 depending on pixel intensity |
| Feedforward inputs burst width | 6 ms |
| No. of AMPA+NMDA inputs per burst | 60 for active context, 0 otherwise |
| Max AMPA+NMDA weight | 0.2 nS |
| Min AMPA+NMDA weight | 0.01 nS |
| Contextual inputs burst width | 20 ms |
| Soma comp AMPA (GABA) background rate | 100 (200) Hz |
| Dend comps AMPA (GABA) background rate | 60 (120) Hz |
| Weight AMPA (GABA) background synapses | 1 (1) nS |

To ensure that the combined feedforward post-synaptic potential (PSP) varied within a reasonable dynamic range, we took the feedforwards weights derived previously (ΔPMD, ΔSD, PCA, and RP to the hidden layer, and 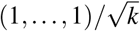 to the output neuron) and applied a scale factor

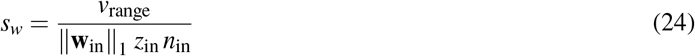

to the input weight vector **w**_in_ of each neuron, with *z*_in_ the somatic input resistance of the L5 PC, *v*_range_ a voltage range parameter and *n*_in_ the number of feedforward inputs (*n* = 784 for a neuron in the hidden layer and *k* = 100 for the output neuron in multitask EMNIST, and *n* = 2 and *k* = {4, 10} for the boolean tasks). *v*_range_ is a parameter that allowed us to fine tune the dynamic range of the combined feedforward PSPs. For the hidden neurons, *v*_range_ was set heuristically to 120 mV on multitask EMNIST and 60 mV on the boolean tasks, and for the output neuron, *v*_range_ was set to 450 mV on multitask EMNIST and 600 mV on the boolean tasks. Together with *z*_in_, this yielded feedforward weights in nA.

To train this system, we presented 600000 samples for each task in multitask EMNIST (balanced across task-classes, so repetition of the same sample may occur) by converting them to Gaussian input spike bursts (the number of input spikes in a burst varied between 0 and 10 and was proportional to pixel intensity), and injecting these burst in the network at 150 ms intervals. We then froze the plasticity rule and tested performance on 500 samples for each task taken from the EMNIST test set (again balanced across task-classes).

### Task-modulated contrastive learning

For TMCL (Fig 6), we convert CIFAR-10^61^ into a multitask learning problem (multitask CIFAR-10) as before, by defining 10 1-vs-all classification tasks, and sampled data in a balanced manner across tasks and task-classes. We kept 9000 samples as validation set. Each color channel was centered, then normalized to unit variance.

We implement a visual geometry group-like (VGG) architecture following Illing et al.^33^ Precisely, we train a stack of *L* convolutional layers with kernel size 3, stride 1 and 64 channels, and applied batch normalization^93^ to the task-modulated convolution outputs before applying the ReLU activation function. Batch normalization renders the task-independent bias *b* obsolete; it was not included in our simulations. By consequence, the output of a convolutional unit was

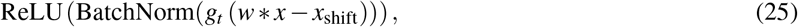

where *x* denotes the image patch, *w* refers to the respective convolutional filter, * denotes the convolution operation, and *g*_*t*_ the task-specific gain. Every second layer was succeeded by a MaxPool layer with stride 2×2.

Our approach learns a stack of convolutional layers iteratively, by adding the next layer on top of the previously learnt ones. Each iteration consists of two distinct phases: first, feedforward filters to the next layer are learned through CL, and subsequently we learn task- and neuron-specific gains for the new layer in a supervised manner.

For CL phase, we follow the SimCLR algorithm^60^. We use the same image augmentations that the authors used on their CIFAR-10 experiments:

~~~
random_resized_crop(32, scale=(0.08, 1.0), ratio=(1., 1.))
random_horizontal_flip(p=0.5)
color_jitter(brightness=0.4, contrast=0.4, saturation=0.4, hue=0.1, p=0.8)
random_grayscale(p=0.2)
~~~

followed by standard score normalization according to the statistics of the original dataset. To train the filters from layer *l* − 1 to layer *l* (*l* = 1, …, *L*), we first generate batches of augmented hidden representations in layer *l* − 1. With *x*_1_, …, *x*_*N*_ a batch of input samples (see Table 4 for batch size), we generate an augmented batch 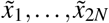 of twice the original size, with samples 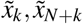 being positive pairs, i.e. generated through image augmentations from the same source sample *x*_*k*_. We then propagate this augmented batch through the hierarchy to layer *l* − 1, while applying task-gains from a random task to each augmented sample, to obtain a batch of task-modulated hidden representations 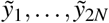. Next, the convolutions with the to be learned filters from layer *l* − 1 to layer *l* are computed and the ReLU is applied to obtain the representation that is fed into the CL-MLP. The CL-MLP consists of a hidden layer and an output layer with dimension 64, and uses ReLU activation. The similarity *s*_*i, j*_ between representations *z*_*i*_ and *z* _*j*_ obtained from final layer of the CL-MLP is computed as their cosine similarity

**Table 4.**
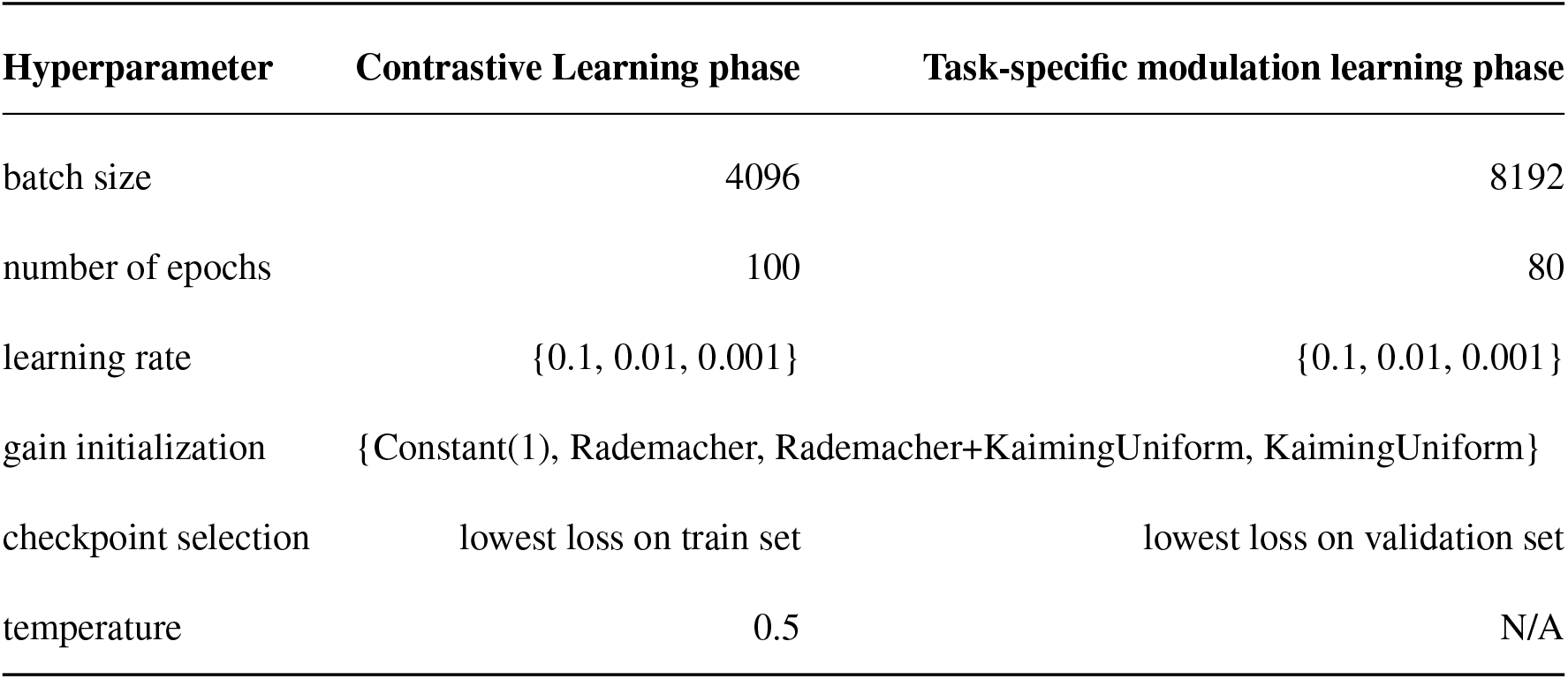
Hyper-parameters used in our simulations. Sets denote that the hyper-parameter is part of a grid search with the given values.

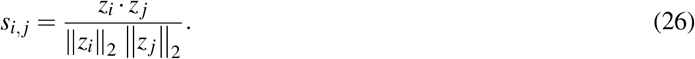

The loss was computed as

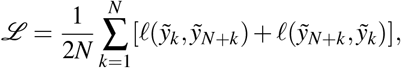

with

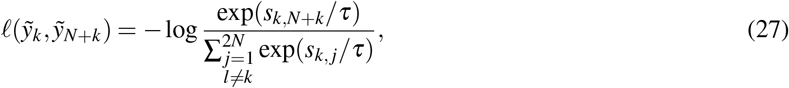

and temperature *τ* = 0.5. The error gradient of this loss is then used to train the filters from layer *l* − 1 to layer *l*. All parameters, including convolutional filters, for layers *< l* − 1 remained frozen. For *l* = 1, layer *l* − 1 is the input layer and no task-modulation could be applied, so that 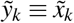.

To perform the classification for all one-vs-all tasks, we reduce the height and width axes by averaging, retaining a 64-dimensional vector with the per-channel averages. Finally, an output unit applies tanh(*·*) to the inner product of the 64-dimensional representation vector and a learned, task-independent weight vector **w**_out_. Task-specific gains in layer *l* are then trained to minimize the classification loss (same as in our fully connected architectures) at the output unit (test performance at this output unit is what is reported in Fig 6B,C), and the process is continued recursively at layer *l* + 1.

Except for the task-specific gains, all parameters (i.e. the CL-MLP parameters, convolutional filters, the x-shift and the output unit) follow the Kaiming initialization, a standard approach in the VGG literature^94^. For the task-specific gains, we tested the following initialization strategies

- **Constant(1):** Initialize each entry to 1.
- **Rademacher:** Sample each entry from Uniform{−1, 1}.
- **KaimingUniform:** Each entry is sampled i.i.d. from Uniform(−*γ, γ*) with 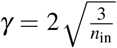 and where *n*_in_ is the number of inputs targeting a given unit.
- **Rademacher+KaimingUniform:** Sample each entry from **Rademacher**, then add a sample from **KaimingUniform** to each entry.

The final performance numbers reported were selected from the grid search described in Table 4, according to the best accuracy on the validation set. All experiments were implemented in PyTorch^89^ and training was performed using Adam^90^.

---

**Figure S1.**
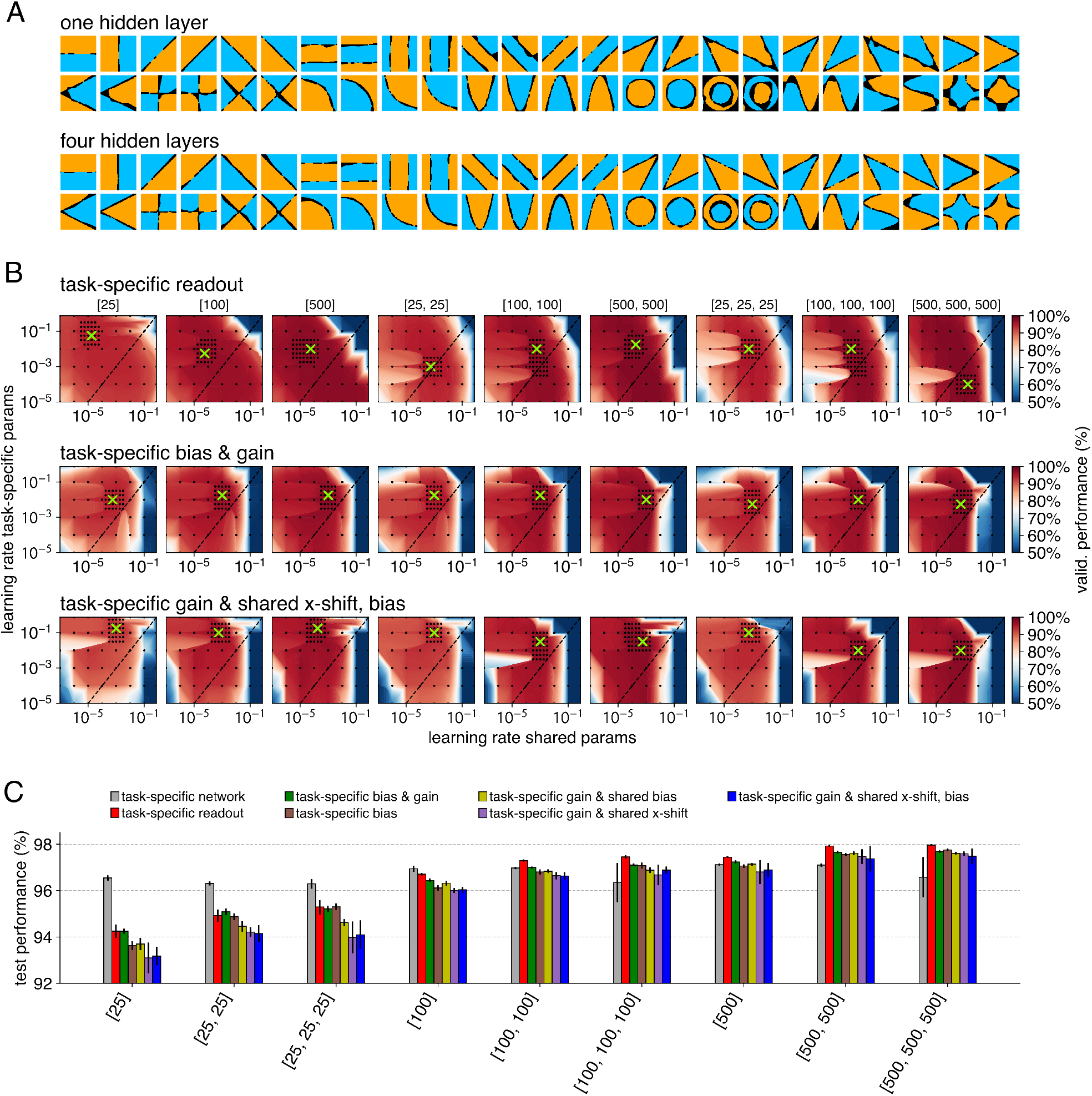
Supplementary results for the networks trained in a fully supervised fashion. **A:** Full set of binary classification tasks (48), solved with a network with a single hidden layer (top) and four hidden layers (bottom) through adaptation of the task-specific gains. Each hidden layer consisted of 50 units. Feedforward weights, x-shifts and biases were shared across tasks. **B:** Performances on multitask EMNIST of task-specific networks (no parameters shared across tasks, grey), task-specific readouts (red) and the various possible neuron-specific modulations for all architectures (x-labels denote the number of hidden units in each layer). **C:** Results of learning rate scans for the three multitask learning approaches shown in Fig 3, for all network architectures. For neuron-specific modulations (middle and bottom rows) the optimal (green cross) modulation learning rate (y-axis) was generally larger than the weight learning rate (x-axis).

**Figure S2.**
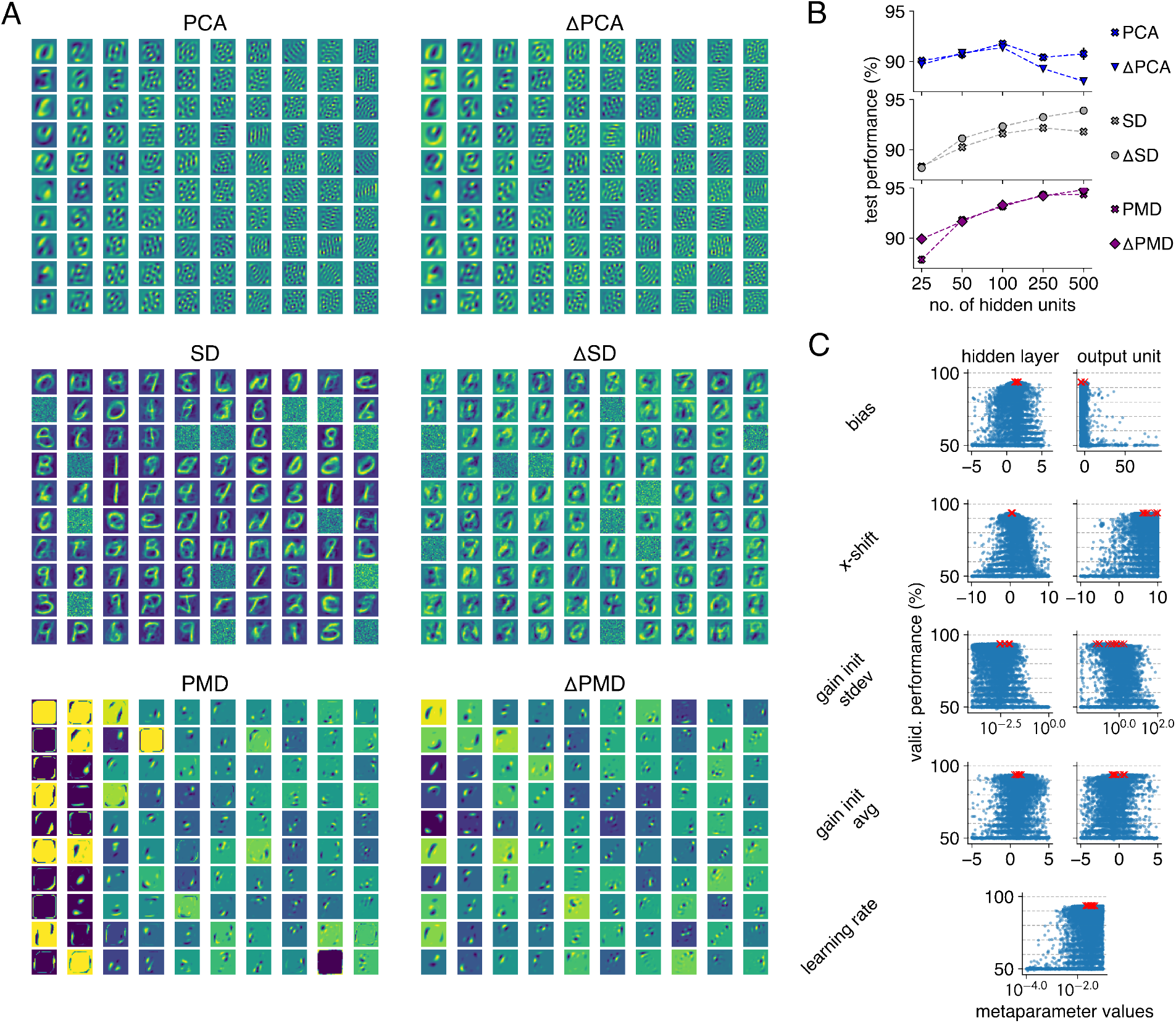
Supplementary results for the networks with unsupervised feedforward weights and supervised gain adaptation. **A:** Weight vectors to the hidden units (*k* = 100) for the three unsupervised learning algorithms Fig 4, i.e. principal component analysis (top), applied to data samples (PCA) or difference vectors (ΔPCA), sparse dictionary learning (middle) applied to data samples (SD) or differences (ΔSD), and penalized matrix decomposition (bottom) applied to data samples (PMD) or differences (ΔPMD). **B:** Comparison of performance of gain-modulated networks on multitask EMNIST, between feedforward weights given by applying the respective unsupervised learning algorithms to the data samples versus the difference vectors, as a function of layer size. For PCA and ΔPCA, weight vectors are very similar (A), resulting in similar performance for *k* ≤ 100. For larger *k*, performance decreases as the extra principal components are no longer useful for classification tasks. For SD and PMD applied to data samples vs. differences, the distinct weight vectors (A) result in a performance increase for ΔSD resp. ΔPMD vs. SD resp. PMD. **C:** To combine the unsupervised weights with supervised gain modulation to learn the specific tasks, we performed an evolutionary meta-parameter optimization on the shared bias, the shared x-shift, the gain learning rate and the gain initialization. Note that shared parameters were not only shared across tasks, but also across neurons. Gain initialization was Gaussian with optimized mean (gain init avg) and standard deviation (gain init stdev). Plots show performance of a randomly chosen subset of ten tasks evaluated on the validation set for ΔPMD with *k* = 100, for all configurations tried by the evolutionary algorithm. The ten best metaparameter configurations are marked by red crosses. The optimal metaparameter configuration was det4e6rmined in this way for each input weight matrix and hidden layer size shown in Fig 4F.

**Figure S3.**
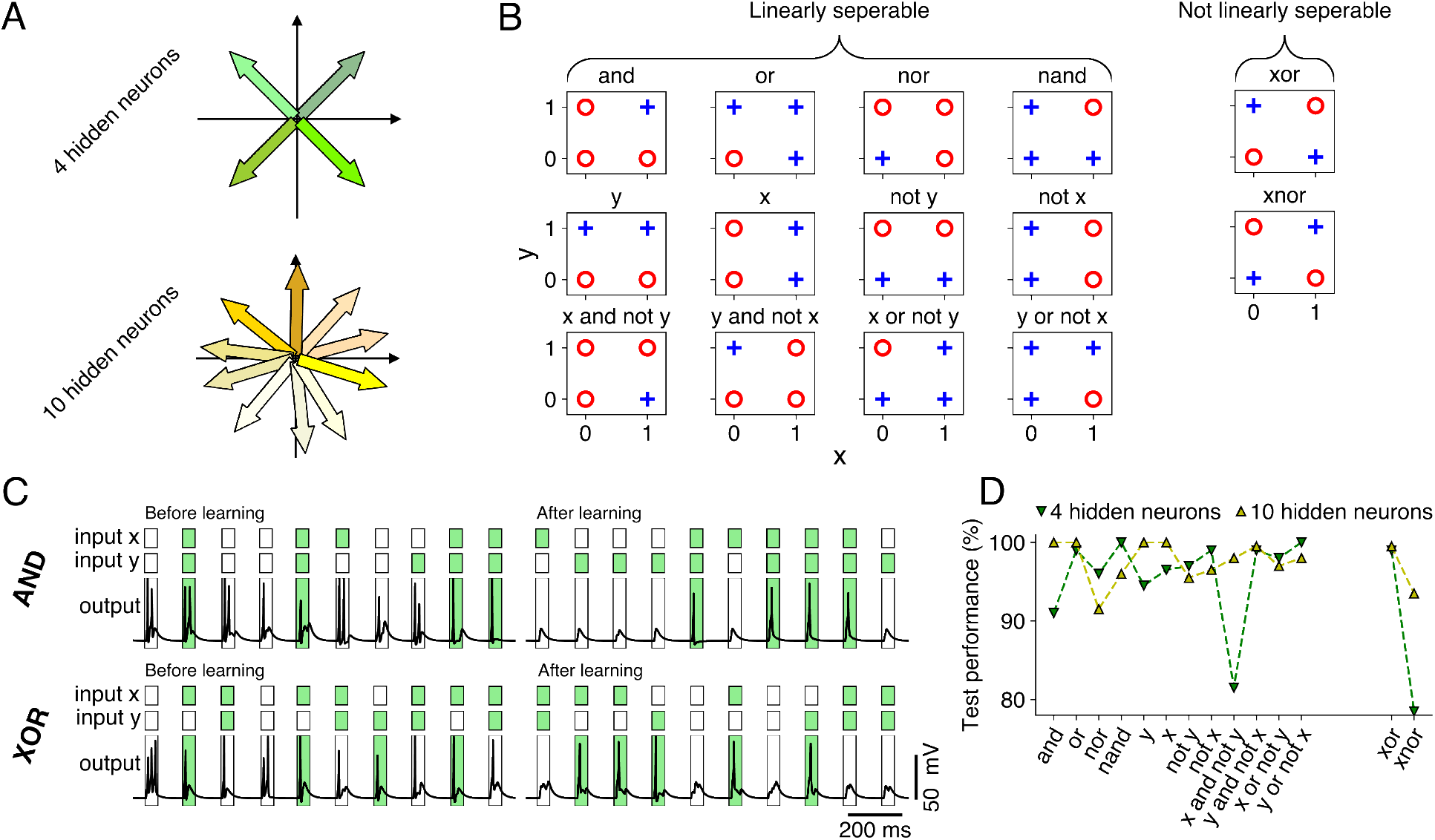
Application of the biophysically realistic spiking network architecture to all non-trivial boolean functions. **A:** Two network configurations were trained, one with 4 hidden neurons and one with 10 hidden neurons. The weight vectors were distributed so as to allow construction of decision boundaries with normal vectors in any direction of the two-dimensional input space. For 4 hidden neurons (top) we chose the centers of the 4 quadrants, whereas for 10 hidden neurons (bottom) we divided the unit circle in 10 equal parts and sampled a vector at random from each of these parts. **B:** We learn the 14 possible non-trivial boolean classification tasks on inputs *x, y* ∈ {0, 1}. 12 of these tasks are linearly separable (left) and two are linearly non-separable (right). The output neuron of the network should learn to spike in response to the blue crosses, and not spike in response to the red circles. **C:** Voltage response of the output neuron (in the network with 10 hidden neurons) to inputs *x, y* ∈ {0, 1}. Note that input values were sampled in such a way that for each task the network received a balanced set of input combinations with + and ° targets. Input values {0, 1} were converted to a Gaussian burst of 6 ms width containing 20 spikes in case of 1, and no spikes otherwise. The output neuron spikes indiscriminately initially (left) but learns to spike correctly after learning (right), here shown for the “and” (top) “xor” (bottom) task. **D**: Performance (best of 5 initialization seeds) for the networks with 4 and 10 hidden neurons.

**Figure S4.**
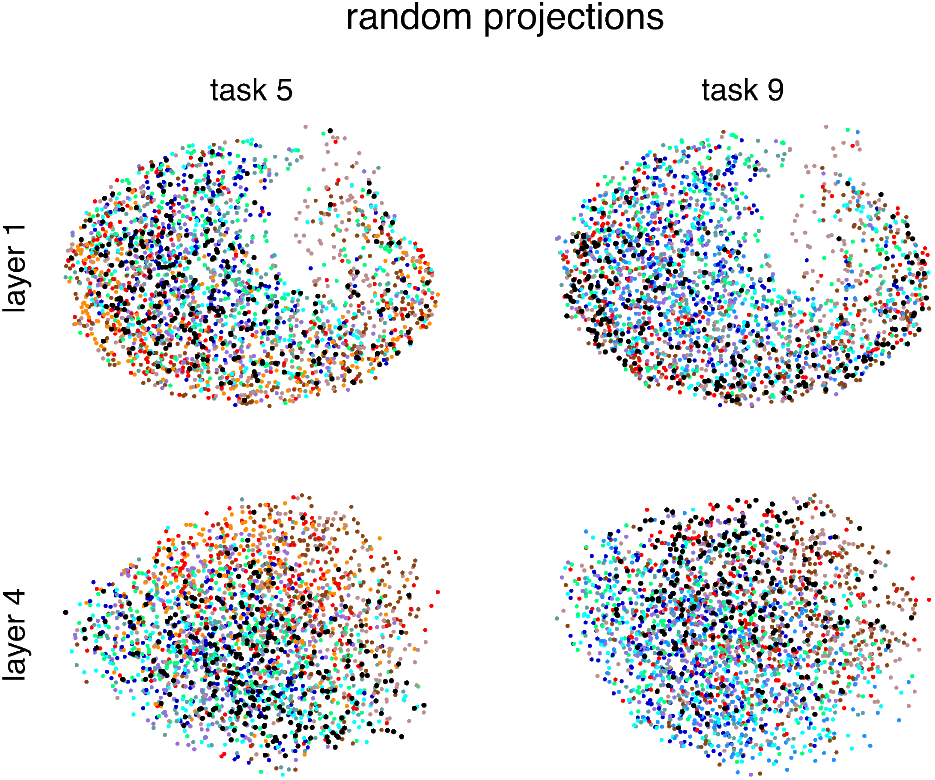
UMAP of the hidden representations, as in Fig 6F, but for the case of RP.

## References

1. Riesenhuber, M. & Poggio, T. Hierarchical models of object recognition in cortex. Nature Neuroscience 2, 1019–1025 (1999).

2. Siegle, J. H. et al. Survey of spiking in the mouse visual system reveals functional hierarchy. Nature (2021).

3. Hubel, D. H. & Wiesel, T. N. Receptive fields, binocular interaction and functional architecture in the cat’s visual cortex. The Journal of physiology 160, 106–154 (1962).

4. Formisano, E. et al. Mirror-Symmetric Tonotopic Maps in Human Primary Auditory Cortex. Neuron 40, 859–869 (2003).

5. Bruce, C., Desimone, R. & Gross, C. G. Visual properties of neurons in a polysensory area in superior temporal sulcus of the macaque. Journal of Neurophysiology 46, 369–384 (1981).

6. Landi, S. M., Viswanathan, P., Serene, S. & Freiwald, W. A. A fast link between face perception and memory in the temporal pole. Science 373, 581–585 (2021).

7. Mesgarani, N. & Chang, E. F. Selective cortical representation of attended speaker in multi-talker speech perception. Nature 485, 233–236 (2012).

8. Chan, A. M. et al. Speech-Specific Tuning of Neurons in Human Superior Temporal Gyrus. Cerebral Cortext 24, 2679–93 (2014).

9. Yamins, D. L. K. et al. Performance-optimized hierarchical models predict neural responses in higher visual cortex. Proceedings of the National Academy of Sciences 111, 8619–8624 (2014).

10. Hong, H., Yamins, D. L. K., Majaj, N. J. & DiCarlo, J. J. Explicit information for category-orthogonal object properties increases along the ventral stream. Nature Neuroscience 19, 613–622 (2016).

11. Gilbert, C. D. & Li, W. Top-down influences on visual processing. Nature Reviews Neuroscience 14, 350–363 (2013).

12. Roth, M. M. et al. Thalamic nuclei convey diverse contextual information to layer 1 of visual cortex. Nature Neuroscience 19, 299–307 (2016).

13. Mineault, P. J., Tring, E., Trachtenberg, J. T. & Ringach, D. L. Enhanced Spatial Resolution During Locomotion and Heightened Attention in Mouse Primary Visual Cortex. The Journal of Neuroscience 36, 6382–6392 (2016).

14. Busse, L. et al. Sensation during Active Behaviors. The Journal of Neuroscience 37, 10826–10834 (2017).

15. Atiani, S., Elhilali, M., David, S. V., Fritz, J. B. & Shamma, S. A. Task Difficulty and Performance Induce Diverse Adaptive Patterns in Gain and Shape of Primary Auditory Cortical Receptive Fields. Neuron 61, 467–480 (2009).

16. Rutten, S., Santoro, R., Hervais-adelman, A., Formisano, E. & Golestani, N. Cortical encoding of speech enhances task-relevant acoustic information. Nature Human Behaviour (2019).

17. Popovkina, D. V. & Pasupathy, A. Task Context Modulates Feature-Selective Responses in Area V4. The Journal of Neuroscience 42, 6408–6423 (2022).

18. Keller, G. B., Bonhoeffer, T. & Hübener, M. Sensorimotor Mismatch Signals in Primary Visual Cortex of the Behaving Mouse. Neuron 74, 809–815 (2012).

19. Pakan, J. M., Currie, S. P., Fischer, L. & Rochefort, N. L. The Impact of Visual Cues, Reward, and Motor Feedback on the Representation of Behaviorally Relevant Spatial Locations in Primary Visual Cortex. Cell Reports 24, 2521–2528 (2018).

20. Banerjee, A. et al. Value-guided remapping of sensory cortex by lateral orbitofrontal cortex. Nature 585, 245–250 (2020).

21. Fiser, A. et al. Experience-dependent spatial expectations in mouse visual cortex. Nature Neuroscience 19, 1658–1664 (2016).

22. Vélez-Fort, M. et al. A Circuit for Integration of Head- and Visual-Motion Signals in Layer 6 of Mouse Primary Visual Cortex. Neuron 98, 179–191.e6 (2018).

23. Doron, G. et al. Perirhinal input to neocortical layer 1 controls learning. Science 370, eaaz3136 (2020).

24. Shin, J. N., Doron, G. & Larkum, M. E. Memories off the top of your head. Science 374, 538–539 (2021).

25. Ruder, S. An Overview of Multi-Task Learning in Deep Neural Networks. arXiv preprint (2017). 1706.05098.

26. Crawshaw, M. Multi-Task Learning with Deep Neural Networks: A Survey (2020). 2009.09796.

27. Cardin, J. A. Functional flexibility in cortical circuits. Current Opinion in Neurobiology 58, 175–180 (2019).

28. Perez, E., Strub, F., De Vries, H., Dumoulin, V. & Courville, A. FiLM: Visual Reasoning with a General Conditioning Layer. Proceedings of the AAAI Conference on Artificial Intelligence 32 (2018).

29. Sun, T. et al. Learning Sparse Sharing Architectures for Multiple Tasks. Proceedings of the AAAI Conference on Artificial Intelligence 34, 8936–8943 (2020).

30. Iyer, A. et al. Avoiding Catastrophe: Active Dendrites Enable Multi-Task Learning in Dynamic Environments. Frontiers in Neurorobotics 16 (2022). 2201.00042.

31. Oja, E. Principal components, minor components, and linear neural networks. Neural Networks 5, 927–935 (1992).

32. Brito, C. S. N. & Gerstner, W. Nonlinear Hebbian Learning as a Unifying Principle in Receptive Field Formation. PLOS Computational Biology 12, e1005070 (2016).

33. Illing, B., Ventura, J., Bellec, G. & Gerstner, W. Local plasticity rules can learn deep representations using self-supervised contrastive predictions. In Advances in Neural Information Processing Systems, vol. 34, 30365–30379 (Curran Associates, Inc., 2021).

34. Perrett, D. I. et al. Organization and functions of cells responsive to faces in the temporal cortex. Philosophical Transactions of the Royal Society of London. Series B: Biological Sciences 335, 23–30 (1992).

35. Thorpe, S., Fize, D. & Marlot, C. Speed of processing in the human visual system. Nature 381, 520–522 (1996).

36. Tremblay, R., Lee, S. & Rudy, B. GABAergic Interneurons in the Neocortex: From Cellular Properties to Circuits. Neuron 91, 260–292 (2016).

37. Chance, F. S., Abbott, L. F. & Reyes, A. D. Gain Modulation from Background Synaptic Input. Neuron 35, 773–782 (2002).

38. Ferguson, K. A. & Cardin, J. A. Mechanisms underlying gain modulation in the cortex. Nature Reviews Neuroscience 21, 80–92 (2020).

39. Self, M. W., Kooijmans, R. N., Supèr, H., Lamme, V. A. & Roelfsema, P. R. Different glutamate receptors convey feedforward and recurrent processing in macaque V1. Proceedings of the National Academy of Sciences 109, 11031–11036 (2012).

40. Manita, S. et al. A Top-Down Cortical Circuit for Accurate Sensory Perception. Neuron 86, 1304–1316 (2015).

41. Schiller, J., Major, G., Koester, H. & Schiller, Y. NMDA spikes in basal dendrites of cortical pyramidal neurons. Nature 1261, 285–289 (2000).

42. Major, G., Larkum, M. E. & Schiller, J. Active properties of neocortical pyramidal neuron dendrites. Annual review of neuroscience 36, 1–24 (2013).

43. Major, G., Polsky, A., Denk, W., Schiller, J. & Tank, D. W. Spatiotemporally Graded NMDA Spike/Plateau Potentials in Basal Dendrites of Neocortical Pyramidal Neurons (Supplementary figures). Journal of neurophysiology (2008).

44. Larkum, M. E. The Guide to Dendritic Spikes of the Mammalian Cortex In Vitro and In Vivo 19 (2022).

45. Hay, E., Hill, S., Schürmann, F., Markram, H. & Segev, I. Models of neocortical layer 5b pyramidal cells capturing a wide range of dendritic and perisomatic active properties. PLoS computational biology 7, e1002107 (2011).

46. Haider, B., Häusser, M. & Carandini, M. Inhibition dominates sensory responses in the awake cortex. Nature 493, 97–100 (2013).

47. Destexhe. Inhibitory “noise”. Frontiers in Cellular Neuroscience (2010).

48. Cohen, G., Afshar, S., Tapson, J. & van Schaik, A. EMNIST: An extension of MNIST to handwritten letters (2017). 1702.05373.

49. Li, N. & DiCarlo, J. J. Unsupervised Natural Experience Rapidly Alters Invariant Object Representation in Visual Cortex. Science 321, 1502–1507 (2008).

50. Marblestone, A. H., Wayne, G. & Kording, K. P. Toward an Integration of Deep Learning and Neuroscience. Frontiers in Computational Neuroscience 10 (2016).

51. Illing, B., Gerstner, W. & Brea, J. Biologically plausible deep learning — But how far can we go with shallow networks? Neural Networks 118, 90–101 (2019). 1905.04101.

52. Olshausen, B. A. & Field, D. J. Code for Natural Images. Nature 381, 607–609 (1996).

53. Mairal, J., Bach, F., Ponce, J. & Sapiro, G. Online dictionary learning for sparse coding. In Proceedings of the 26th Annual International Conference on Machine Learning - ICML ‘09, 1–8 (ACM Press, Montreal, Quebec, Canada, 2009).

54. Witten, D. M., Tibshirani, R. & Hastie, T. A penalized matrix decomposition, with applications to sparse principal components and canonical correlation analysis. Biostatistics (Oxford, England) 10, 515–534 (2009).

55. Wybo, W. A. et al. Data-driven reduction of dendritic morphologies with preserved dendro-somatic responses. eLife 10, e60936 (2021).

56. Rumelhart, D. E., Hintont, G. E. & Williams, R. J. Learning representations by back-propagating errors 4 (1986).

57. Goodfellow, I., Bengio, Y. & Courville, A. Deep Learning (The MIT Press, 2016).

58. Lillicrap, T. P., Santoro, A., Marris, L., Akerman, C. J. & Hinton, G. Backpropagation and the brain. Nature Reviews Neuroscience 21, 335–346 (2020).

59. LeCun, Y., Kavukcuoglu, K. & Farabet, C. Convolutional networks and applications in vision. In Proceedings of 2010 IEEE International Symposium on Circuits and Systems, 253–256 (IEEE, Paris, France, 2010).

60. Chen, T., Kornblith, S., Norouzi, M. & Hinton, G. A Simple Framework for Contrastive Learning of Visual Representations. In Proceedings of the 37th International Conference on Machine Learning, 1597–1607 (PMLR, 2020).

61. Krizhevsky, A. Learning Multiple Layers of Features from Tiny Images. Technical Report TR-2009, University of Toronto, Toronto (2009).

62. McInnes, L., Healy, J. & Melville, J. UMAP: Uniform Manifold Approximation and Projection for Dimension Reduction (2020). 1802.03426.

63. Ziv, Y. et al. Long-term dynamics of CA1 hippocampal place codes. Nature Neuroscience 16, 264–266 (2013).

64. Driscoll, L. N., Pettit, N. L., Minderer, M., Chettih, S. N. & Harvey, C. D. Dynamic Reorganization of Neuronal Activity Patterns in Parietal Cortex. Cell 170, 986–999.e16 (2017).

65. Deitch, D., Rubin, A. & Ziv, Y. Representational drift in the mouse visual cortex. Current Biology 31, 4327–4339.e6 (2021).

66. Schoonover, C. E., Ohashi, S. N., Axel, R. & Fink, A. J. P. Representational drift in primary olfactory cortex. Nature 594, 541–546 (2021).

67. Marks, T. D. & Goard, M. J. Stimulus-dependent representational drift in primary visual cortex. Nature Communications 12, 5169 (2021).

68. Kalle Kossio, Y. F., Goedeke, S., Klos, C. & Memmesheimer, R.-M. Drifting assemblies for persistent memory: Neuron transitions and unsupervised compensation. Proceedings of the National Academy of Sciences 118, e2023832118 (2021).

69. Rule, M. E. & O’Leary, T. Self-healing codes: How stable neural populations can track continually reconfiguring neural representations. Proceedings of the National Academy of Sciences 119, e2106692119 (2022).

70. Bosman, C. A. et al. Attentional Stimulus Selection through Selective Synchronization between Monkey Visual Areas. Neuron 75, 875–888 (2012).

71. van Kerkoerle, T. et al. Alpha and gamma oscillations characterize feedback and feedforward processing in monkey visual cortex. Proceedings of the National Academy of Sciences 111, 14332–14341 (2014).

72. Bastos, A. M. et al. Visual Areas Exert Feedforward and Feedback Influences through Distinct Frequency Channels. Neuron 85, 390–401 (2015).

73. Michalareas, G. et al. Alpha-Beta and Gamma Rhythms Subserve Feedback and Feedforward Influences among Human Visual Cortical Areas. Neuron 89, 384–397 (2016).

74. Richter, C. G., Thompson, W. H., Bosman, C. A. & Fries, P. Top-Down Beta Enhances Bottom-Up Gamma. The Journal of Neuroscience 37, 6698–6711 (2017).

75. Richter, C. G., Coppola, R. & Bressler, S. L. Top-down beta oscillatory signaling conveys behavioral context in early visual cortex. Scientific Reports 8, 6991 (2018).

76. Poirazi, P., Brannon, T. & Mel, B. W. Pyramidal neuron as two-layer neural network. Neuron 37, 989–999 (2003).

77. Hawkins, J. & Ahmad, S. Why Neurons Have Thousands of Synapses, a Theory of Sequence Memory in Neocortex. Frontiers in Neural Circuits 10, 1–13 (2016). 1511.00083.

78. Kawaguchi, Y. Pyramidal Cell Subtypes and Their Synaptic Connections in Layer 5 of Rat Frontal Cortex. Cerebral Cortex 27, 5755–5771 (2017).

79. Bannister, A. P. Inter- and intra-laminar connections of pyramidal cells in the neocortex. Neuroscience Research 53, 95–103 (2005).

80. Schuman, B., Dellal, S., Prönneke, A., Machold, R. & Rudy, B. Neocortical Layer 1: An Elegant Solution to Top-Down and Bottom-Up Integration. Annual Review of Neuroscience 44, 221–252 (2021).

81. Larkum, M. E., Zhu, J. J. & Sakmann, B. A new cellular mechanism for coupling inputs arriving at different cortical layers. Nature 398, 338–41 (1999).

82. Larkum, M. E., Senn, W. & Lüscher, H.-R. Top-down Dendritic Input Increases the Gain of Layer 5 Pyramidal Neurons. Cerebral Cortex 14, 1059–1070 (2004).

83. Doron, M., Chindemi, G., Muller, E., Markram, H. & Segev, I. Timed Synaptic Inhibition Shapes NMDA Spikes, Influencing Local Dendritic Processing and Global I/O Properties of Cortical Neurons. Cell Reports 21, 1550–1561 (2017).

84. Wybo, W. A., Torben-Nielsen, B., Nevian, T. & Gewaltig, M. O. Electrical Compartmentalization in Neurons. Cell Reports 26, 1759–1773.e7 (2019).

85. Carnevale, N. T. & Hines, M. L. The NEURON Book (2004).

86. Rotter, S. & Diesmann, M. Exact digital simulation of time-invariant linear systems with applications to neuronal modeling. Biological cybernetics 81, 381–402 (1999).

87. Jahr, C. E. & Stevens, C. F. A quantitative description of NMDA receptor-channel kinetic behavior. The Journal of neuroscience : the official journal of the Society for Neuroscience 10, 1830–1837 (1990).

88. Behabadi, B. F. & Mel, B. W. Mechanisms underlying subunit independence in pyramidal neuron dendrites. Proceedings of the National Academy of Sciences of the United States of America 111, 498–503 (2014).

89. Paszke, A. et al. PyTorch: An Imperative Style, High-Performance Deep Learning Library. In Advances in Neural Information Processing Systems 32, 8024–8035 (2019).

90. Kingma, D. P. & Ba, J. Adam: A Method for Stochastic Optimization. In Proceedings of the 3rd International Conference for Learning Representations (arXiv, 2015). 1412.6980.

91. Pedregosa, F. et al. Scikit-learn: Machine Learning in Python. Journal of Machine Learning Research 12, 2825–2830 (2011).

92. Fortin, F.-A., De Rainville, F.-M., Gardner, M.-A., Parizeau, M. & Gagné, C. DEAP: Evolutionary Algorithms Made Easy. Journal of Machine Learning Research 13, 2171–2175 (2012).

93. Ioffe, S. & Szegedy, C. Batch Normalization: Accelerating Deep Network Training by Reducing Internal Covariate Shift. In Proceedings of the 32nd International Conference on Machine Learning (JMLR.org, Lille, 2015). 1502.03167.

94. He, K., Zhang, X., Ren, S. & Sun, J. Delving deep into rectifiers: Surpassing human-level performance on imagenet classification. Proceedings of the IEEE International Conference on Computer Vision 2015 Inter, 1026–1034 (2015). 1502.01852v1.

